# Three-dimensional mapping of human lung fibrosis identifies spatial coupling between tissue microarchitecture and maladaptive repair states

**DOI:** 10.64898/2026.11.05.764175

**Authors:** Joseph A. Bell, Stephanie K. Robinson, Robert Pedley, Gautam Ahuja, Adam Szmul, Ian Reddin, Jaya Thomas, Jeanette Norman, Sonya James, Philip J. Basford, Victor Barozi, Dipanjan Bhattacharya, James G. H. Parkin, Rebekah Allen, Lucy N. Sayer, Jon Ward, Grace Haydon, Nina Vyas, Sanjay Jogai, David Johnston, Aiman Alzetani, Serena J. Chee, Christian H. Ottensmeier, Ben G. Marshall, Tim Wallis, Sumeet Mahajan, Gareth Thomas, Emily Offer, Gayle Marshall, Kerstin Geillinger-Kästle, Peter Seither, Matthew Thomas, Andres F. Vallejo, Michael Eyres, Yihua Wang, Donna E. Davies, Joseph Jacob, Aurelie Fabre, Orestis L. Katsamenis, Kedar Natarajan, Mark G. Jones

## Abstract

Human lung fibrosis progressively destroys three-dimensional tissue architecture, yet how the structural microenvironment shapes cellular states remains unclear. Integrating 3D X-ray histology with spatial omics, we generate a volumetric multimodal atlas of human lung fibrosis that enables direct interrogation of relationships between tissue microarchitecture and spatial molecular programmes. We identify collapse of the alveolar-capillary unit - the coupled loss of alveolar type 1 epithelial cells, capillary endothelial cells, and basement membrane integrity - as a structural transition associated with early fibrogenesis. Surrounding transitional zones exhibit aberrant epithelial differentiation and proteolytic matrix remodelling, with epithelium shifting toward bronchiolised, airway-like states. Stromal activation converges on spatially restricted fibrogenic cores, sites of maximal collagen maturation and matrix stabilisation that function as signalling hubs within surrounding perifocal and vascular microenvironments. Molecular signatures of atlas-defined tissue states were associated with lung function and mortality across independent patient cohorts. Together, these data support a model in which fibrosis emerges as a spatial tissue-state process, with alveolar-capillary collapse redirecting repair towards mechanically stabilised niches that reinforce pathological remodelling.

## Introduction

Lung fibrosis is a progressive and fatal disorder characterised by aberrant remodelling of the distal lung parenchyma. Despite approved antifibrotic therapies, median survival remains only 3-5 years, reflecting the absence of interventions that effectively halt or reverse disease. In healthy lungs, epithelial injury typically triggers regenerative repair, whereas in lung fibrosis this process instead leads to maladaptive remodelling and deposition of aberrant extracellular matrix (ECM).^1^

The lung is an elastic organ with a complex three-dimensional (3D) branching structure. It comprises a network of airways that conduct air from the trachea to millions of thin-walled alveoli, where gas exchange occurs across the alveolar-capillary membrane.^2^ Fibrotic remodelling disrupts this architecture in spatially heterogeneous ways, resulting in regions of alveolar loss, extracellular matrix remodelling, the replacement of alveolar architecture by airway-like (bronchiolised) structures, and vascular distortion.^3^ These structural changes are increasingly recognised to arise in spatially organised multicellular niches.^4–6^ Spatial-profiling studies have identified dysregulated cell states and epithelial-stromal-immune interactions in human lung fibrosis.^4–14^ For example, Vannan et al. used subcellular-resolution spatial transcriptomics across more than one million cells to define discrete epithelial-stromal-immune niches and proposed a trajectory from alveolar dysregulation to advanced fibrosis.^14^ Despite these advances, current spatial approaches largely infer tissue organisation from two-dimensional sections, limiting our ability to relate molecular cell states to the three-dimensional tissue architecture that underpins progressive fibrosis.

To begin to address this challenge, we previously established a 3D X-ray histology (XRH) platform enabling non-destructive micro-computed tomography of intact formalin-fixed, paraffin-embedded (FFPE) tissue blocks.^15–18^ This approach generates high-fidelity micron-scale maps of tissue microarchitecture.^15,16,19–24^

Here, we describe a multimodal methodology that integrates spatial transcriptomic and proteomic profiling with high-resolution XRH volumes, enabling molecular and cellular states to be interpreted within intact 3D lung architecture. The resulting volumetric atlas reconstructs a spatial trajectory of human lung fibrosis, linking quantitative features of tissue microarchitecture to changes in cellular composition, regulation and function. These findings support a model in which structural failure and maladaptive cellular states are spatially coupled, reinforcing maladaptive repair and progressive fibrosis.

## Results

### Quantitative 3D X-ray histology resolves heterogeneous microarchitectural remodelling

An overview of the study is provided in Figure 1A. High-resolution XRH (4.9-8.7 µm voxels) was applied to diagnostic surgical lung biopsy FFPE tissue from individuals with idiopathic pulmonary fibrosis (IPF) and non-fibrotic controls, enabling three-dimensional analysis of the structural microenvironment during active fibrogenesis at the time of diagnosis. Virtual XRH sections closely matched histological sections following alignment to the sectioning plane (Fig. S1A-C), confirming morphological fidelity of XRH (Fig. 1B-C). XRH volumes revealed marked three-dimensional structural heterogeneity in IPF, including loss of normal alveolar geometry and regions of increased tissue mass density associated with architectural distortion (Fig. 1D-E).

**Fig. 1:**
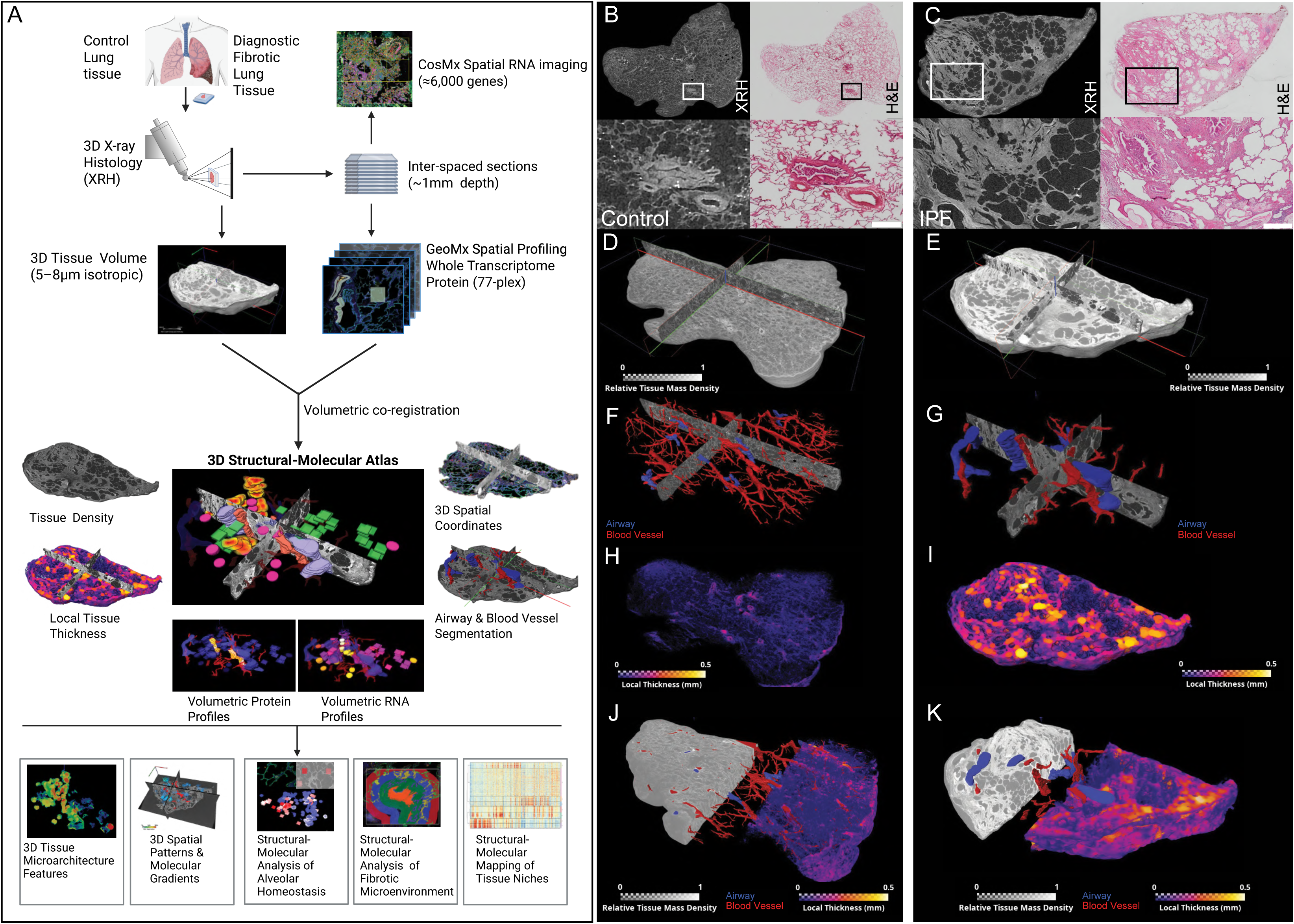
3D X-ray histology (XRH) identifies complex anatomical heterogeneity in fibrotic lung tissue. **(A)** Study overview. Formalin-fixed paraffin embedded lung tissue blocks from control and IPF donors underwent X-Ray Histology (XRH) imaging to capture intact 3D tissue architecture, followed by sectioning for histology and spatial RNA and protein profiling. Histological sections and spatial data were co-registered into XRH volumes. Schematic created with BioRender.com **(B, C)** Co-registered XRH virtual sections and H&E sections for control (B) and IPF (C) tissue; insets show higher-magnification view. Scale bars, 500µm. **(D, E)** 3D-XRH reconstructions visualising surface-rendered tissue mass density with orthogonal XY and XZ/YZ planes for control (**D**) and IPF (**E**) volumes. **(F, G)** Airway (blue) and blood vessel (red) segmentations in control (**F**) and IPF volumes (**G**). (**H**, **I**) Local tissue-thickness mapping projected onto the XRH volume for control (**H**) and IPF (**I**). **(J-K)** Composite 3D-rendering of density, airway, blood vessels and tissue thickness for control (**J**) and IPF (**K**).

We next performed machine-learning-assisted segmentation of airways and blood vessels to generate three-dimensional coordinate maps, enabling quantitative assessment of spatial organisation and providing a spatial reference framework for subsequent integration with molecular datasets (Fig. 1F-G). In control lungs, vessels formed coherent, hierarchical networks, whereas in IPF, vascular architecture was disrupted, with vascular structures frequently truncated within fibrotic regions. Local tissue-thickness mapping further highlighted spatially heterogeneous thickening of alveolar structures in IPF, in contrast to the uniformly thin septal architecture in controls (Fig. 1H-I). Quantitative XRH metrics (Fig. S1D-E) confirmed elevated tissue mass density and increased local tissue thickness in IPF. Together, these volumetric analyses established the three-dimensional structural framework for spatial molecular integration (Fig. 1J-K).

### Multiscale spatial omics maps cellular and molecular niches across anatomical compartments

Following XRH scanning, inter-spaced tissue sections (≈100µm apart; ten-per-block) underwent Bruker GeoMx whole-transcriptome profiling (≈18,000 genes), with an adjacent section assayed using a 77-plex protein panel. Regions of interest (ROIs) were selected across distinct compartments, including alveolar septae, native bronchovascular airway walls, vascular walls, and immune-rich (CD45^+^) aggregates (Fig. 2A & Fig. S2A-B). To identify discrete stromal regions of active extracellular-matrix remodelling, we used tenascin-C (TNC), a matricellular glycoprotein induced during tissue injury and repair and implicated in fibrotic niche formation.^25,26^ Analysis of our previous digital spatial profiling^4^ and laser-capture microdissection RNA-seq^9^ datasets demonstrated that fibroblast foci, sites of active lung fibrogenesis, were encriched for *TNC* and interstitial collagens (e.g. *COL1A2*), whereas *ACTA2* (α-SMA) was broadly expressed across mesenchymal and vascular compartments (Fig. S2C,D). Immunofluorescence confirmed focal TNC protein enrichment, in contrast to the widespread ACTA2 expression (Fig. S2E-H), supporting TNC as a marker of spatially restricted fibrogenic regions (TNC^+^ fibrotic cores).

**Fig. 2:**
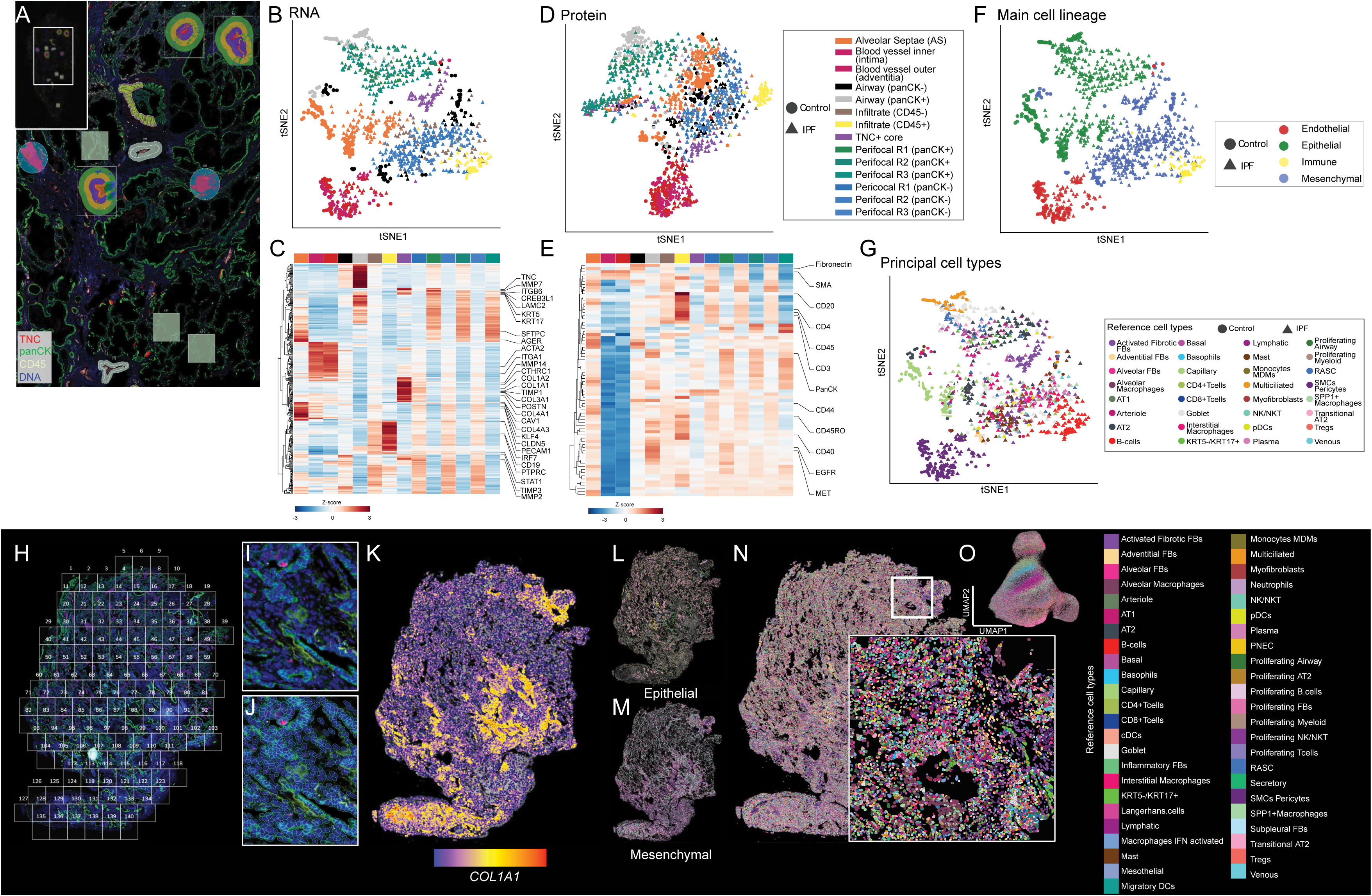
Multimodal spatial profiling resolves epithelial, stromal, vascular and immune niches. Inter-spaced tissue sections (≈100µm apart; ten-per-block) underwent spatial transcriptomic (GeoMx whole transcriptome ~18,000 genes; n=1,257 ROIs) or proteomic (77-plex; n=1,362 ROIs) profiling **(A)** Regions-of-interest (ROI) across alveolar septae, vascular wall (intima and adventitia), airway wall, CD45^+^ immune-rich aggregates, and TNC^+^ cores with three concentric perifocal regions (R1-R3) extending into the surrounding microenvironment. Selected ROIs were computationally segmented into PanCK^+^/PanCK^−^ (epithelial/stromal) or CD45^+^/CD45^−^ (immune/non-immune) compartments. **(B)** t-SNE of transcriptome profiles with ROI coloured by anatomical compartment. **(C)** Hierarchical clustering of region-specific differentially expressed genes **(D)** t-SNE of spatial proteomic profiles with ROI coloured by anatomical compartment. **(E)** Spatial protein heatmap **(F, G)** t-SNE coloured by (**F**) main cell lineages and (**G**) deconvolution-inferred principal cell type. (**H-O**) Single-cell resolution spatial transcriptomics (CosMx; ~6,000 genes) was performed on one representative tissue section from each tissue block, generating a dataset comprising 944,439 cells containing 2.97 × 10^8^ spatial transcripts. Panels H-N show representative images. (**H**) Representative IPF tissue section with analysed fields of view. (**I,J**) Immunofluorescence morphology markers used for cell segmentation, showing tissue morphology (**I**) and the corresponding segmentation mask (**J**). (**K**-**N**) Spatial maps showing (K) COL1A1 transcript expression, (**L**) epithelial cell types, (**M**) mesenchymal cell types, and (**N**) annotated cell types. (**O**) UMAP embedding of all CosMx-profiled cells coloured by annotated cell type. In (**C**) and (**E**), colour scale represents z-score normalised expression or abundance across all ROIs.

Using TNC^+^ fibrotic cores as spatial anchors, we sampled the surrounding microenvironment across three concentric perifocal regions (Perifocal R1-3), each 90 µm thick and extending ~270 µm from the TNC^+^ boundary (Fig. S2B,I). ROIs were further segmented by PanCK or CD45 expression where appropriate, generating paired epithelial/stromal or immune/parenchymal profiles within the same anatomical region (Fig. S2B,I).

After quality control, 1257 RNA ROIs and 1362 protein ROIs were retained for analysis (Fig. S2A). Dimensional reduction of transcriptomic profiles revealed clear segregation by disease status, anatomical context, and compartment-specific marker expression (Fig. 2B-C, Fig. S2J, Table S1A). Protein profiling using an immune-focussed panel similarly resolved ROI classes, with vascular regions, CD45^+^ immune aggregates, and alveolar septae forming distinct clusters (Fig. 2D-E).

Cellular composition was inferred using reference cell identities from a published spatial lung fibrosis atlas (>1.5 million cells).^14^ This identified distinct epithelial, mesenchymal, endothelial and immune populations across anatomical niches (Fig. 2F), including multiciliated epithelial cells within PanCK^+^ airways and activated fibrotic fibroblasts within TNC^+^ cores (Fig. 2G, S2K). Independent lineage-signature deconvolution confirmed AT1 and AT2 populations within alveolar septae (Fig. S2L).

To complement regional GeoMx profiling we generated subcellular spatial transcriptomics using CosMx (≈6,000 genes) on selected sections (Fig. 2H-O). Image analysis, segmentation and quality control yielded 944,439 cells containing 2.97×10^8^ spatial transcripts (Fig. 2H-J). Mapping of the fibrillar collagen gene *COL1A1* demonstrated spatially restricted and heterogeneous expression (Fig. 2K). Reference-based cell annotation corroborated niche-specific cellular composition and spatial segregation of epithelial, stromal, endothelial and immune populations (Fig. 2L-O), with high *COL1A1* transcript expression localising predominantly to mesenchymal-rich areas (Fig. 2K,M).

### Co-registration of 3D microarchitecture and spatial molecular profiles establishes a structurally anchored volumetric atlas

To integrate molecular programmes with 3D lung architecture, spatial omics ROIs were co-registered to XRH volumes and extended into 100 µm-thick volumes of interest (VOIs; Fig. 3A-D). This generated 1,257 RNA VOIs and 1,362 protein VOIs (Fig. 3A-D), each integrating 3D spatial coordinates with matched transcriptomic, proteomic and structural features including tissue mass density, septal thickness, and geodesic distance to the nearest vessel or airway (Fig. 3E-H).

**Fig. 3:**
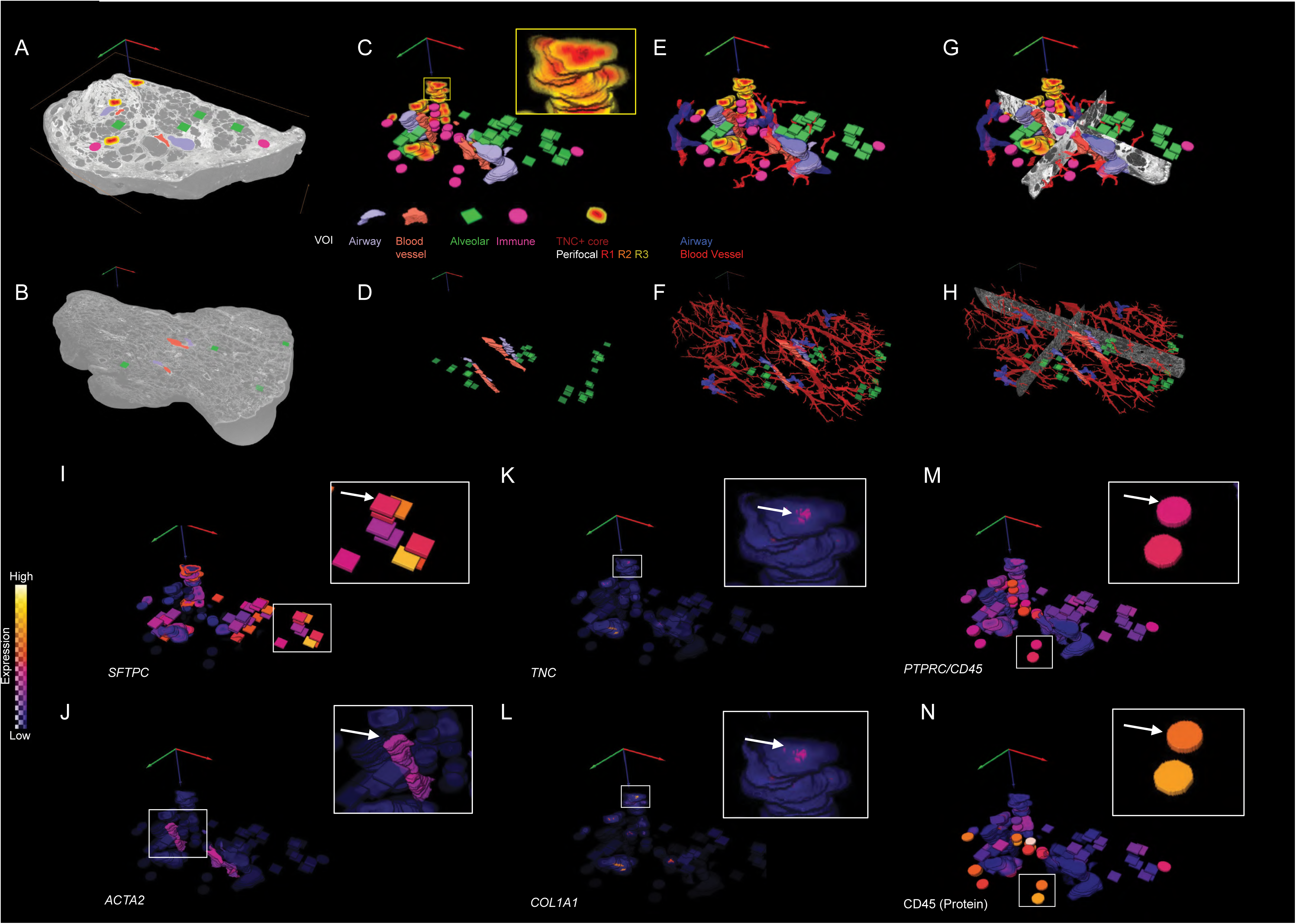
**3D integration of X-ray histology and spatial molecular profiling**. **(A,B)** Surface-rendered 3D XRH volumes for IPF (A) and control (B) tissue with volumes of interest (VOIs) overlaid (one plane visualised for clarity). **(C,D)** Three-dimensional visualisation of VOIs in IPF (C) and control (D) tissue volumes, coloured by VOI type. Inset in (C) shows higher-magnification view of a TNC^+^ core with concentric perifocal regions. **(E,F)** VOIs overlaid on segmented airway (blue) and vascular (red) structures within the 3D tissue volume **(G,H)** Fully integrated atlas combining VOIs, segmented airway and vascular structures, and orthogonal XRH slices for control (G) and IPF (H). **(I-N)** VOI-level gene and protein expression within IPF XRH volume, coloured by molecular abundance: *SFTPC* (I), *ACTA2* (J), *TNC* (K), *COL1A1* (L), *PTPRC* (M), and CD45 protein (N). White-bordered insets show higher-magnification views of representative high-expression VOI clusters, with arrows indicating corresponding zoomed region.

Across the VOI classes, projection of canonical marker genes confirmed their anatomical identity (Fig. 3I-N, S2J). The alveolar epithelial marker *SFTPC* was enriched within alveolar septae (Fig. 3I); *ACTA2* delineated smooth-muscle rich vascular walls (Fig. 3J); *TNC* together with *COL1A1* marked discrete stromal zones of active fibrogenesis (Fig. 3K-L); and *PTPRC* (CD45) identified immune aggregates (Fig. 3M), corresponding to CD45 protein localisation (Fig. 3N) in three-dimensional space.

The resulting atlas is provided as an open-access resource, available for download and interactive exploration through MoBIE, a Fiji plugin (Fig. S3A-D).^27,28^

### Tissue mass density defines structural trajectories of alveolar-capillary unit collapse

A striking feature of the XRH volumes was a marked but heterogeneous increase in mass density of alveolar septae in IPF compared with control tissue (Fig. 4A-C). These high-density alveolar regions were accompanied by local thickening of the alveolar walls and architectural distortion. We therefore asked whether increasing septal density was associated with changes in the cellular and molecular composition of the alveolar-capillary gas-exchange unit.

**Fig. 4:**
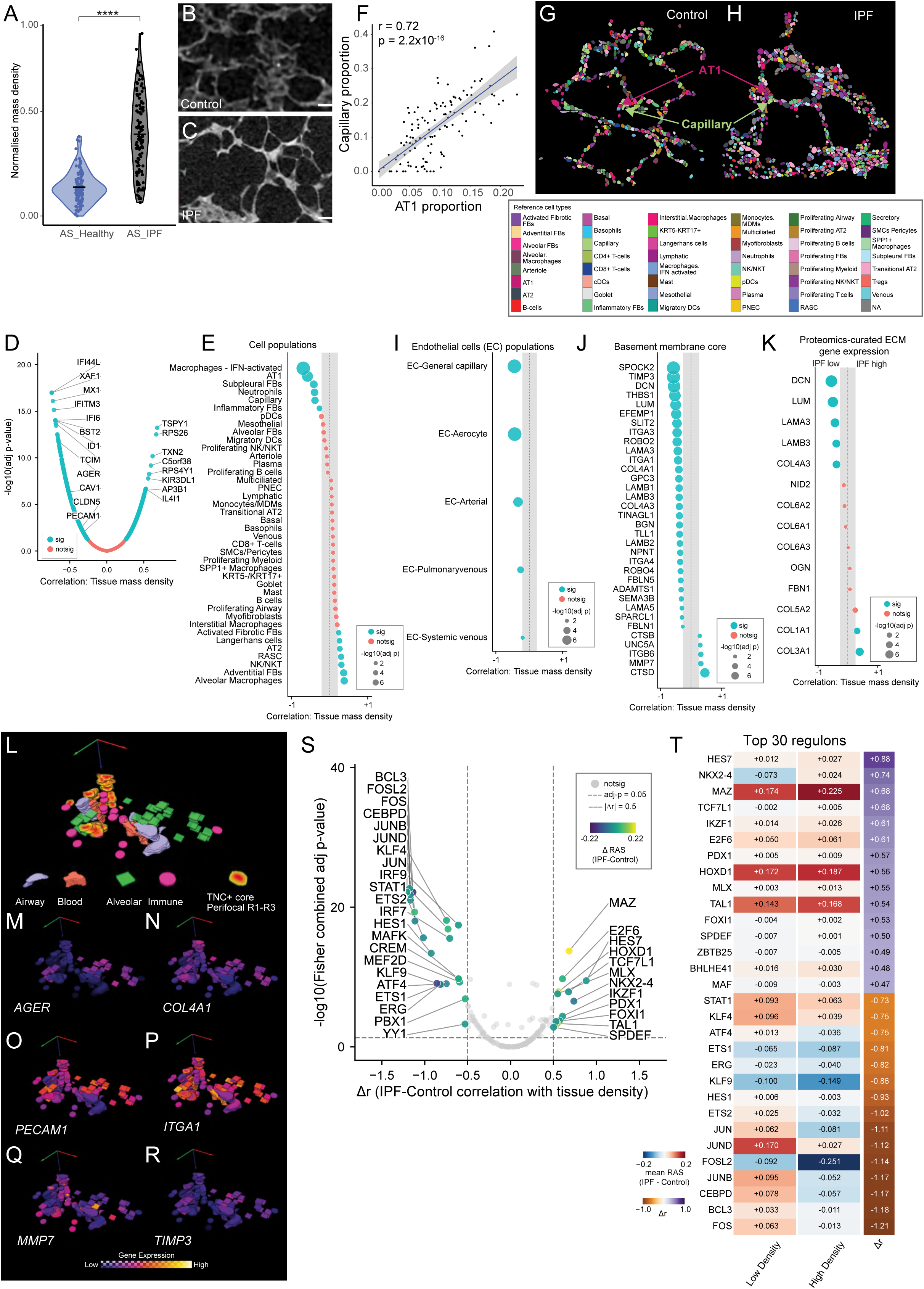
Collapse of the alveolar-capillary exchange unit accompanies increasing alveolar septal density. **(A)** XRH-derived tissue mass density across control and IPF alveolar septae (****p < 0.0001, t-test). **(B, C)** Representative XRH virtual sections of control (B) and IPF (C) alveolar septae. **(D)** Volcano plot showing differential gene expression and its correlation (Spearman r) with tissue mass density in IPF septae. **(E)** Deconvolution-inferred cell-type proportions and their correlation (Spearman r) with tissue mass density across IPF septae. **(F)** Pairwise correlation between AT1 and capillary endothelial cell proportions. **(G,H)** Representative CosMx single-cell spatial cell-type maps in control (**G**) and IPF (**H**) alveolar septae. **(I)** Gene set variation analysis (GSVA) scores for endothelial subtype gene sets and correlation with tissue mass density. **(J)** Correlation of top 30 basement membrane genes (BMbase) with tissue mass density. **(K)** Correlation of gene expression with tissue mass density in IPF tissue, using a curated ECM gene list identified from an independent quantitative human lung proteomic dataset of ECM proteins as differentially abundant (IPF high or IPF low) in IPF compared to control tissue.^30^ **(L-R)** 3D visualisation of VOI types within an IPF tissue volume, coloured by VOI type (L) with 3D XRH overlaid gene expression of (**M**) *AGER*, (**N**) *COL4A1*, (**O**) *PECAM1*, (**P**) *ITGA1*, (**Q**) *MMP7*, and (**R**) *TIMP3*. **(S)** Volcano plot highlighting differential regulons across IPF and control septae, alongside their correlation to tissue density. Each point indicates a differential mean Regulon Activity Score (ΔRAS; IPF-Control) with tissue-density correlation Δr (IPF-Control, x-axis) and regulon significance (-log10 Fisher combined adjusted p-value, y-axis). Dashed lines for significance (adj p=0.05; | Δr |= 0.5). **(T)** Heatmap of top 30 differentially correlated regulons stratified across low- and high-density ROIs (normalised median density=0.206), with mean differential regulon activity scores (ΔRAS, IPF-Control) and differential regulon-density correlation (Δr, rightmost column).

Across IPF alveolar septae, increasing tissue mass density was strongly associated with progressive loss of structural markers of the gas-exchange interface. AT1 epithelial genes (*AGER*, *CAV1*) showed significant negative correlations with density, alongside depletion of endothelial markers (*PECAM1*, *CLDN5*) (Fig. 4D, Table S2A). Cellular deconvolution confirmed these transcriptional changes: AT1 cell proportions declined with increasing density (Fig. 4E) and closely tracked loss of capillary cells (Fig. 4F; r=0.72, p=2.2×10^−16^), consistent with co-ordinated loss of epithelial and vascular compartments rather than selective depletion of a single lineage. In contrast, AT2 epithelial cells increased with density, together with respiratory secretory epithelial cells, adventitial and activated fibrotic fibroblast signatures (Fig. 4E). These compositional shifts were confirmed using our CosMx single-cell spatial transcriptomic maps, which demonstrated reduced AT1 and capillary cell frequency (Fig. 4G-H, S4A). Endothelial subtype analysis identified aerocyte (aCap) and general capillary (gCap) endothelial cell types as the principal vascular populations declining with increasing density (Fig. 4I).

Because epithelial and endothelial cells are anchored by a laminin-rich basement membrane, we next examined expression of canonical basement-membrane genes.^29^ Across IPF alveolar septae, these genes showed significant negative correlations with tissue mass density, indicating loss of the laminin-collagen IV basement-membrane network as septal architecture collapsed (Fig. 4J).

These density-associated basement-membrane changes were further supported by analysis of an extracellular-matrix enriched quantitative proteomic dataset comparing IPF and control tissue.^30^ Proteins reported to be reduced in IPF showed progressively lower gene expression with increasing tissue density, whereas proteins increased in IPF showed the opposite pattern (Fig. 4K). Representative basement membrane components, including *COL4A3*, exhibited negative associations with density, whereas fibrillar collagens including *COL1A1* showed a positive association, consistent with replacement of basement-membrane scaffolds by interstitial matrix. A similar directional relationship was observed using a separate ECM-focussed proteomic dataset (Fig. S4B),^31^ whilst direct comparison of alveolar septae from IPF and control tissue demonstrated depletion of basement-membrane genes and enrichment of fibrillar collagen genes within IPF septae (Fig. S4C).

Septae also exhibited evidence of reprogramming of matrix turnover and adhesion with increasing tissue density. Among matrix-remodelling enzymes, *MMP7*, an epithelial-derived protease and established circulating biomarker associated with disease severity and outcome in IPF^32,33^ was the strongest positive correlate of septal density, while the matrix-bound inhibitor *TIMP3* was markedly reduced (Fig. 4J), indicating a shift toward pericellular proteolysis. In parallel, integrins mediating epithelial-endothelial basement-membrane anchorage (*ITGA1*, *ITGA3*) declined with increasing density, whereas the epithelial wound-associated integrin *ITGB6* increased.^34^

3D visualisation of gene expression illustrated substantial spatial heterogeneity in the localisation of epithelial (*AGER*), basement-membrane (*COL4A1*), endothelial (*PECAM1*) and adhesion (*ITGA1*) markers across alveolar regions (Fig. 4L-P, S4G). In contrast, *MMP7* and *TIMP3* exhibited largely anticorrelated spatial distributions (Fig. 4Q,R, Fig. S4G), suggesting the emergence of a protease-dominant microenvironment within high-density septae.

At the pathway level, comparison of control vs IPF alveolar septae regions corroborated these observations, with downregulation of angiogenesis and cell adhesion programmes within IPF alveolar septae (Fig. S4D). Collagen cross-linking and fibril-organisation programmes were not enriched, indicating that high-density septae are characterised by matrix disorganisation rather than formation of a mature fibrotic matrix.

To further define regulatory programmes associated with septal collapse, we performed transcription factor regulatory network analysis across alveolar septae and then compared inferred regulatory programmes between high- and low-density regions (Fig. 4S-T, Fig. S4E, H & Table S2B-C). Increased septal density was associated with reduced activity of regulons linked to vascular-epithelial homeostasis (KLF4-, BCL3-, CEBPD), interferon signalling (STAT1, IRF7, IRF9) and AP-1 family mediated injury responses (Fig. 4S-T; S4E-F, H).

In contrast, high-density septae demonstrated enrichment of developmental and lineage-associated regulons, including HES7, MAZ, TCF7L1, IKZF1 and HOXD1 (Fig. 4S-T, S4F,H). Regulons associated with mucociliary differentiation and bronchiolisation-associated epithelial reprogramming, including SPDEF and FOXI1, also increased with septal density (Fig. 4T, S4E). Together, these findings support reactivation of developmental and airway-lineage transcriptional programmes accompanying maladaptive epithelial remodelling within dense septal regions.

In parallel, regulatory activity of the NOTCH effector HES1, implicated in alveolar progenitor maintenance,^35^ declined with increasing tissue density, indicating impaired NOTCH-mediated epithelial repair signalling within collapsing septae (Fig. 4S-T, S4E-H). These regulatory shifts parallel the depletion of AT1 epithelial and capillary endothelial populations and accompany the emergence of maladaptive bronchiolisation-associated epithelial states and broader reorganisation of endothelial-associated programmes across the collapsing alveolar-capillary interface.

Together, these data demonstrate that increasing alveolar septal density is associated with coordinated failure of the epithelial, endothelial and basement-membrane components of the alveolar-capillary exchange unit. This transition is accompanied by suppression of innate immune and homeostatic transcriptional programmes alongside activation of developmental and epithelial lineage-remodelling pathways that define an axis of early fibrogenic injury.

### Vascular remodelling is spatially coupled to surrounding tissue states

To determine whether structural remodelling extended beyond the alveolar compartment in fibrotic lung tissue, we next examined vascular, airway, and immune regions. Blood vessels within IPF tissue exhibited increased tissue mass density (Fig. 5A-C). We spatially profiled the inner and outer halves of arterial blood vessel walls, capturing endothelial-enriched intimal regions (BVi) and stromal-enriched adventitial compartments (BVa) (Fig. S2A-B). Gene-set enrichment analysis identified suppression of vascular development, cell-matrix adhesion, and smooth-muscle contractile programmes across vascular compartments (Fig. 5D-E and Table S3A-B).

**Fig. 5:**
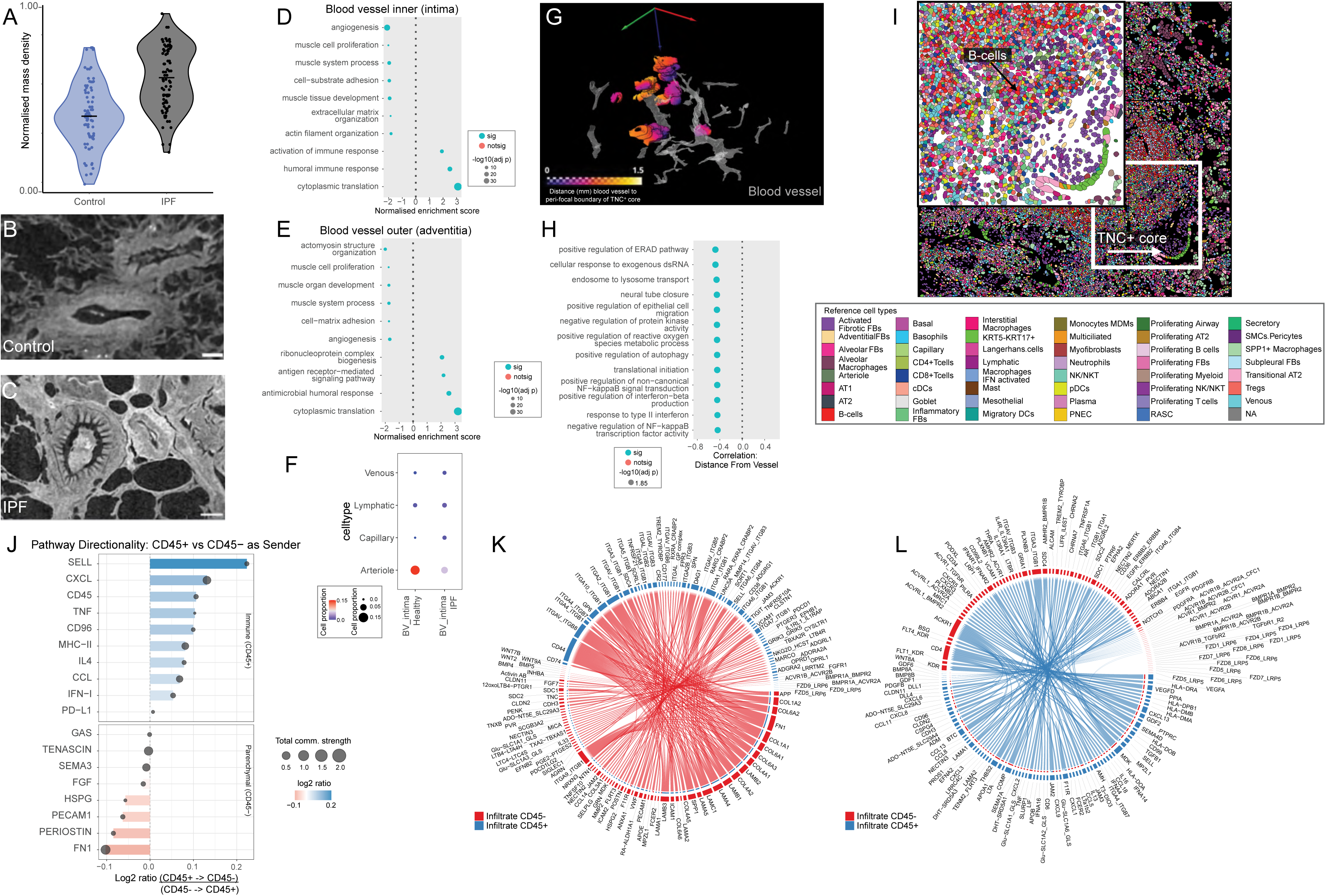
Vascular remodelling follows spatial gradients in fibrotic lung tissue. (A-C) Tissue density distribution across vascular VOI, with representative XRH virtual sections through control **(B)** and IPF **(C)** blood vessels. **(D-E)** Gene set enrichment analysis (GSEA) of biological processes enriched in IPF intimal **(D)** and adventitial **(E)** vascular compartments. **(F)** Endothelial cell subtype proportions (deconvolution) within the intima compartment. (**G**) 3D rendering of segmented blood vessels and perifocal microenvironments surrounding TNC^+^ cores in an IPF tissue block; colour scale indicates geodesic distance from the perifocal boundary to the nearest blood vessel. (**H**) Gene Ontology terms whose pathway-level enrichment scores correlate with vessel-to-perifocal-boundary distance, ranked by Spearman correlation; negative correlations indicate progressive loss of pathway activity with increasing distance from vasculature. **(I)** Representative CosMx single-cell spatial cell-type map of a tissue region adjacent to a TNC^+^ core, with the low-magnification indicating the core (white arrow) and high-magnification view showing surrounding cellular architecture, including immune populations. **(J)** Pathway directionality analysis quantifying the relative contribution of CD45^+^ and CD45^−^ compartments as signal senders across curated ligand-receptor pathways. For each pathway, x-axis indicates log₂ ratio CD45^+^-to-CD45^−^ versus CD45^−^-to-CD45^+^ cells total outgoing communication probability; positive values (blue) indicate CD45^+^-dominant sending and negative values (red) indicate CD45^−^-dominant sending. Dot size is proportional to total cross-compartment communication strength. (**K, L**) Chord plots of significant (p<0.05) ligand-receptor interactions between CD45^+^ and CD45^−^ cell subpopulations; CD45^+^-dominant outgoing communications (**K**) and CD45^−^-dominant outgoing communications (**L**). Outer arcs represent individual cell subpopulations; chord width is proportional to interaction strength.

Within the intima, endothelial homeostatic and angiogenic pathways were reduced alongside marked enrichment of immune, ribosome biogenesis, and cytoplasmic translation programmes (Fig. 5D, Table S3A). Similar translational and immune pathway activation was also observed within the adventitia (Fig. 5E, Table S3B). Cellular deconvolution confirmed reduced arteriole-associated endothelial populations within the intima (Fig. 5F), and decreased smooth-muscle cells in the adventitia (Fig. S5A). Regulatory network analysis identified increased MAZ, SMAD4 and STAT1 activity within IPF vascular regions (Fig. S5B), supporting transcriptional reprogramming accompanying vascular remodelling.

At the protein level, IPF vascular compartments exhibited activation of MAPK-ERK/JNK and PI3K-AKT signalling together with enrichment of stress-responsive and immune-associated markers (Fig. S5C-D). Collectively, these findings indicate loss of vascular structural and contractile identity within fibrotic lung tissue.

We next asked whether there was interplay between vascular architecture and surrounding tissue state. Examining gene-expression changes as a function of the three-dimensional distance between the perifocal boundary surrounding each TNC^+^ core and the nearest blood vessel (Fig. 5G) identified spatially organised transcriptional programmes associated with vascular proximity (Fig. 5H, Table S3C). The most significantly associated pathways declined progressively with increasing distance from the vascular network and included programmes linked to proteostasis, vesicular trafficking, interferon responses and cellular stress adaptation. Collectively, these observations suggest that vascular proximity and perifocal fibro-inflammatory states remain spatially coupled within tissue surrounding TNC^+^ fibrogenic cores. In contrast, native bronchovascular airway compartments exhibited comparatively modest molecular changes (Fig. S5E-G and Table S3G-H), and three-dimensional structural analyses did not identify a consistent relationship between proximity to these airways and fibrogenic transcriptional programmes.

We next compared paired CD45^+^ immune-rich and CD45^−^ parenchymal compartments adjacent to TNC^+^ cores (Fig. 5I-L, Fig. S2A-B, S5H-N, Table S3D-E, I-K). CD45^+^ regions were characterised by adaptive immune signatures (CD40, CD8A, CD19) and encirhment of B-cell activation and immune-response pathways, whereas CD45^−^ regions showed extracellular-matrix and cell-adhesion programmes (COL1A1, ACTA2, CTHRC1) (Fig. S5H). Regulatory network analysis identified enrichment of immune regulators (IRF8, STAT6, RUNX3) in CD45^+^ regions and signalling/metabolic regulators (KLF4, STAT3, XBP1, RXRA) in CD45^−^ compartments (Fig. S5I). Protein and single cell spatial profiling corroborated B-cell-enriched immune niches adjacent to TNC^+^ cores (Fig. 5I, S5K)

Ligand-receptor analysis showed that CD45^−^ parenchymal compartments generated substantially more unique outgoing interactions than CD45^+^ regions, dominated by extracellular-matrix pathways including FN1 and periostin (Fig. 5J-L, S5L,N; Table S3E-F,I-K). In contrast, CD45^+^ niches sent regulatory and inflammatory signals, including SELL, IFN-I and PD-L1, defining distinct structural-remodelling and immune-signalling networks. Whereas FN1 represented the dominant signalling network by overall interaction strength and was generated almost exclusively by CD45^−^ parenchymal cells, SELL exhibited the greatest degree of polarised immune-to-parenchymal directionality (Fig. 5K-L, Fig. S5N & Table S3E-F). Together, these findings further support the presence of spatially organised and functionally distinct immune and structural communication networks within fibrotic tissue.

Together, these findings demonstrate that vascular remodelling, immune organisation and fibro-inflammatory signalling remain spatially coupled within fibrotic lung tissue. These observations indicate that local tissue states are structured by spatial proximity to both vascular networks and sites of active fibrogenesis. We therefore next examined transcriptional programmes within concentric perifocal microenvironments surrounding TNC^+^ fibrogenic cores.

### Perifocal transitional niches orchestrate failed epithelial-mesenchymal crosstalk

Using TNC^+^ cores, which identify foci of active stromal matrix synthesis (Fig. 3K,L; Fig. S2A-B, J), as spatial anchors, we examined epithelial and stromal transitions across the surrounding perifocal microenvironment (Fig. 6A). Integrating transcriptional, regulatory and protein-level analyses resolved coordinated changes across the perifocal gradient towards the ECM-producing core (Fig. 6B-C, S6A & Table S4A-B).

**Fig. 6:**
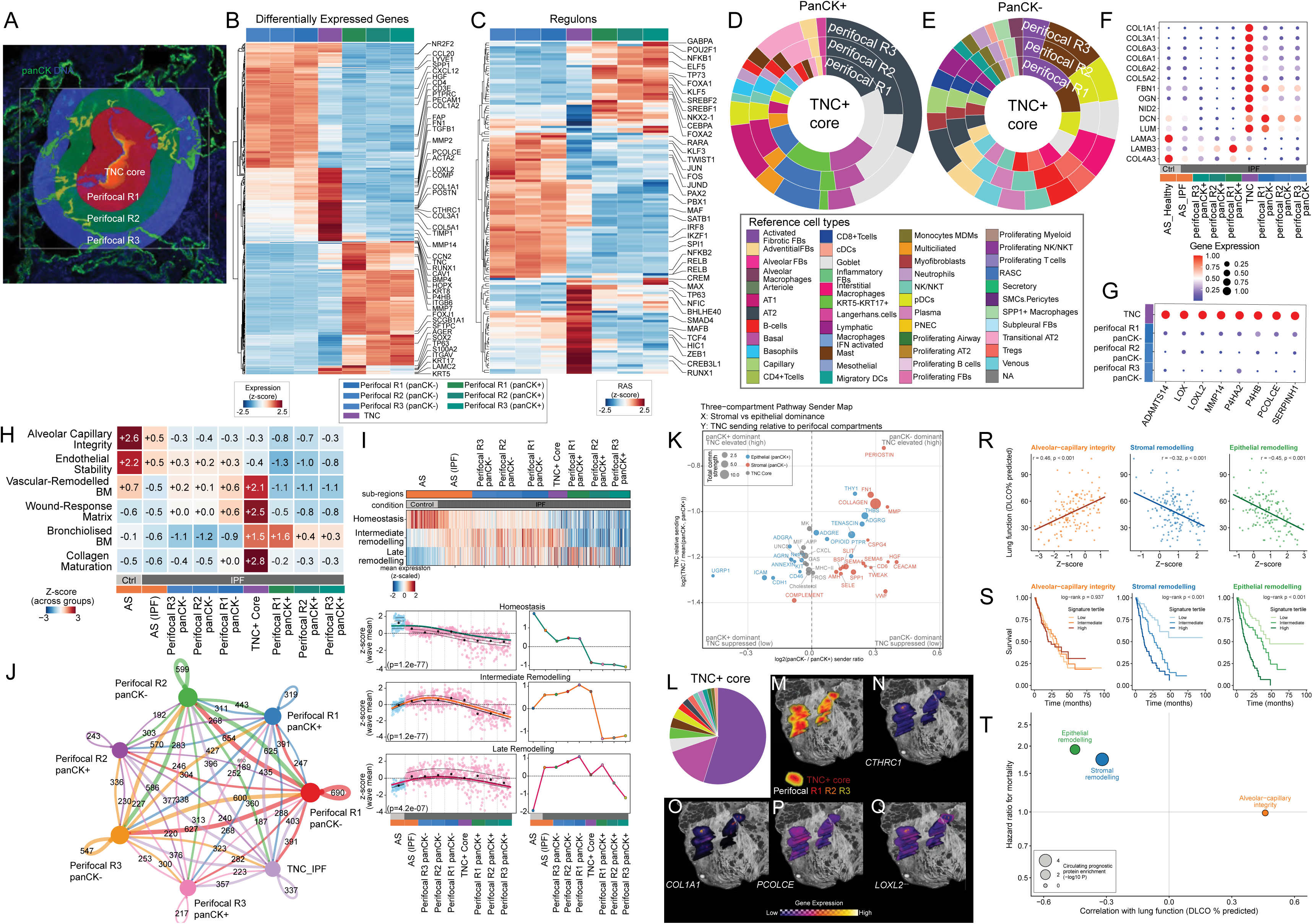
Matrix-stabilised TNC^+^ cores act as signalling hubs for fibrogenic microenvironments. **(A)** Representative GeoMx immunofluorescence showing DAPI (nuclei) and PanCK (green) staining within a captured ROI (white box). TNC^+^ core surrounding concentric perifocal regions (perifocal R1-3) and epithelial (PanCK^+^) and stromal (PanCK^−^) segmented regions indicated. **(B, C)** Hierarchical clustering of differentially expressed genes (**B**) and mean regulon activity scores (**C**) across perifocal compartments and TNC^+^ core. **(D, E)** Cellular composition radial plots for PanCK^+^ epithelial (**D**) and PanCK^−^ stromal regions (**E**). **(F)** Expression of extracellular-matrix proteins across alveolar septae, perifocal regions, and TNC^+^ core, using a curated ECM gene list identified from an independent quantitative human lung proteomic dataset of ECM proteins as differentially abundant in IPF compared to control tissue.^30^ **(G)** Expression of collagen maturation genes across TNC^+^ core and PanCK^−^ perifocal R1-R3 **(H)** Gene-set enrichment across ROI compartments for alveolar-capillary integrity, endothelial stability, vascular-remodelled basement membrane, wound-response matrix, bronchiolised basement membrane, and collagen maturation. Values represent group-level mean enrichment; colour denotes z-score across groups. **(I) Top:** Mean z-scaled expression of alveolar-homeostatic (76 genes), intermediate-remodelling (30 genes) and late-remodelling (93 genes) gene sets defined by Vannan et al.^14^ across individual ROIs. ROIs are ordered and coloured by subregion, scaled tissue density and condition. **Bottom left:** Generalised additive model (GAM) trends for each gene set across individual ROIs. Points represent ROIs coloured by condition; black points indicate z-scaled mean values. **Bottom right:** Corresponding GAM trends calculated using pseudobulk mean values. **(J)** Circle plot of significant ligand-receptor interactions between perifocal compartments and TNC^+^ core (colored nodes). Self-loops denote intra-compartment signalling; edge widths proportional to number of significant interactions. **(K)** Three-compartment pathway sender map showing directional communication dominance across panCK^−^ stromal, panCK^+^ epithelial regions and TNC^+^ core for curated ligand-receptor pathways. X-axis shows log₂ ratio of total outgoing communication probability (PanCK^−^/panCK^+^) and y-axis the log₂ ratio of TNC^+^ core outgoing probability relative to the mean of the two perifocal regions. Quadrant corner annotations summarise directional interpretation (*i.e.* combinatorial epithelial vs stromal dominance alongside TNC interaction signal strengths), dot size is proportional to total cross-compartment communication strength, and colors indicate epithelial (blue), stromal (red), and TNC^+^ core-associated pathways (grey). **(L)** Radial plot showing the inferred cellular composition of the TNC^+^ core. **(M-Q)** Three-dimensional XRH rendering of TNC^+^ cores and surrounding perifocal regions **(M)**, showing gene expression of *CTHRC1* (**N**), *COL1A1* (**O**), *PCOLCE* (**P**), and *LOXL2* **(Q)** within TNC^+^ cores and surrounding perifocal regions. (**R**) Correlation of atlas-derived spatial gene signatures with lung function. Signature activity (singScore, z) plotted against diffusing capacity (DLCO, % predicted) in IPF lung tissue (GSE47460, n = 145)^38^: alveolar–capillary integrity (orange), perifocal stromal remodelling (blue), and perifocal epithelial remodelling (green). Line, linear fit; Spearman r and p values inset. (**S**) Survival by atlas-derived spatial gene signature. Kaplan–Meier curves in bronchoalveolar lavage (GSE70866; n = 176) stratified by signature tertile (low, intermediate and high; light to dark); p, log-rank. Alveolar–capillary integrity (orange), perifocal stromal remodelling (blue), and perifocal epithelial remodelling (green). (**T**) Clinical associations of atlas-derived spatial gene signatures in independent IPF cohorts. Each point represents an atlas-derived spatial gene signature. The x-axis shows the Spearman correlation between signature score and lung function (DLCO, % predicted) in IPF lung tissue (GSE47460), with positive values indicating association with preserved gas transfer. The y-axis shows the GAP-adjusted hazard ratio for mortality per standard deviation of signature score, pooled across three bronchoalveolar lavage cohorts (GSE70866; dashed line, HR = 1). Point size represents enrichment significance (-log₁₀ hypergeometric P value) for proteins encoded by each signature among circulating proteins associated with adverse transplant-free survival in an independent IPF plasma proteomic study.^40^ Green, perifocal epithelial remodelling signature; blue, perifocal stromal remodelling signature; orange, alveolar-capillary integrity signature.

Within the perifocal PanCK^+^ epithelium, differential expression analysis between distal (R3) and core-adjacent (R1) regions revealed progressive loss of alveolar identity and acquisition of aberrant epithelial programmes. Core-adjacent epithelial regions showed enrichment of basaloid-associated markers (*KRT5*, *KRT17*, *S100A2*) and the epithelial remodelling-associated laminin subunit *LAMC2* (Fig. 6B & Table S4A), a component of laminin-332 that is characteristic of airway and basal-cell basement membranes rather than alveolar epithelium.^36^ This transition was accompanied by adhesion reprogramming, with increased expression of integrin subunits such as *ITGAV*. Protein analysis identified receptor tyrosine kinase enrichment in core-adjacent regions, including EGFR and MET, supporting active epithelial remodelling and altered epithelial-matrix engagement (Fig. S6A & Table S4F).

Transcription factor activity analysis identified coordinated loss of homeostatic lung regulators (NKX2-1, CEBPA, FOXA2) alongside increased activity of TP63, SMAD4, and BHLHE40, indicating a basaloid, TGF-β-responsive epithelial state (Fig. 6C & Table S4B). ELF5, which has been implicated in maintaining alveolar epithelial identity^37^, showed the greatest loss of regulatory activity across the perifocal PanCK^+^ gradient (Fig. 6C & Table S4B), suggesting a potential role in gating the transition away from alveolar programmes. In addition, core-adjacent epithelial regions exhibited increased proteolytic activity, with *MMP7* among the most strongly induced transcripts (Fig. 6B), suggesting local basement-membrane disruption.

The perifocal PanCK^−^ stromal compartment exhibited a spatial gradient of extracellular matrix dysregulation converging on the TNC^+^ core. Core-adjacent stromal regions exhibited increased expression of fibrillar collagens (*COL1A1*, *COL3A1*), periostin (*POSTN*), and matrix remodelling genes including *MMP14* (Fig. 6B & Table S4A). Regulatory network analysis identified activation of AP-1 family transcription factors alongside TWIST1 (Fig. 6C & Table S4B), consistent with progressive fibroblast activation and matrix remodelling. RUNX1, a transcription factor associated with mesenchymal cell-state reprogramming, also showed elevated regulon activity in core-adjacent stromal regions (Fig. 6C).

Changes in cellular composition mirrored these transcriptional trajectories (Fig. 6D-E; Fig. S6B & Table S4C). Within the perifocal epithelium, proximity to the core was associated with increased abundance of KRT5^−^/KRT17^+^ cells and marked depletion of AT1 and AT2 populations (Fig. 6D). In the PanCK^−^ stromal compartment, this gradient was accompanied by a transition from alveolar fibroblast populations toward activated fibrotic fibroblasts (Fig. 6E). Immune cell signatures were also evident within PanCK^−^ perifocal regions, but occurred alongside marked expansion of activated fibrotic fibroblast populations. Differential protein analysis demonstrated increased abundance of immune markers in core-adjacent PanCK^+^ regions, including GZMB, CD45, CD3, and CD4, supporting increased immune infiltration and immune-stromal interactions proximal to the core (Table S4F).

In parallel, endothelial composition changed in a compartment-specific manner. Whereas IPF alveolar septae exhibited loss of capillary endothelial signatures relative to control tissue, perifocal PanCK^−^ regions showed emergence of venous and lymphatic endothelial populations not identified within alveolar septae (Fig. 6E, S6B). Together with the depletion of arteriole-associated endothelial programmes in blood vessels, these findings indicate broad spatial re-patterning of endothelial identity across the fibrotic lung microenvironment. This endothelial reorganisation occurred alongside epithelial bronchiolisation in adjacent PanCK^+^ compartments, suggesting coordinated but maladaptive restructuring of epithelial and vascular identities within perifocal niches.

Together, these findings identify perifocal regions as transitional niches characterised by bronchiolisation-associated epithelial remodelling, endothelial-state reorganisation, extracellular matrix remodelling, and progressive fibroblast activation, supporting a model of maladaptive epithelial-mesenchymal cross-talk.

### Matrix stabilisation within the TNC^+^ core as a culmination of failed repair

To determine whether TNC^+^ cores represent a qualitative shift in matrix state, we mapped across the septal-to-core axis the same proteomics-curated ECM genes that showed concordant density-associated expression in IPF alveolar septae (Fig. 4K). Genes encoding fibrillar and provisional matrix components progressively increased through perifocal regions, reaching maximal expression in TNC^+^ cores, whereas homeostatic basement-membrane genes progressively decreased (Fig. 6F-G).

This reciprocal pattern prompted a broader interrogation of gene-set programmes capturing distinct structural, reparative, and stabilisation states of the lung extracellular environment (Fig. 6H, S6C-D). Programmes associated with alveolar-capillary integrity and endothelial stability were already markedly reduced in IPF alveolar septae compared with control septae, indicating early loss of structural support within the alveolar-capillary exchange unit (Fig. 6H). Reactive matrix remodelling programmes exhibited compartment-specific engagement: bronchiolised basement-membrane signatures were most strongly enriched within epithelial perifocal regions adjacent to the TNC^+^ core (Fig. 6H), whereas wound-response matrix and collagen-maturation programmes showed increasing enrichment across perifocal regions, peaking within the TNC^+^ core (Fig. 6H, S6C-D), with significantly greater enrichment in the core relative to all surrounding perifocal regions. Together, these programmes paralleled the transcriptional regulatory shifts observed in perifocal regions (Fig. 6C), linking matrix remodelling to coordinated activation of matrix-producing stromal states and loss of alveolar epithelial homeostatic regulation.

To determine whether our anatomically defined findings aligned with an independently derived remodelling trajectory, we scored each anatomical region for homeostatic, intermediate and late-remodelling gene programmes recently defined by Vannan et al. from computational spatial ordering of alveolar remodelling states across 35 subjects using a targeted image-based spatial transcriptomic panel.^14^ The homeostatic programme declined progressively from alveolar septa towards the TNC^+^ core, whereas intermediate- and late-remodelling programmes were enriched in core-adjacent perifocal regions, particularly within the PanCK^−^ stromal compartment (Fig. 6I). PanCK^+^ perifocal regions exhibited substantially greater inter-ROI transcriptional variability than stromal regions, reflecting a heterogeneous epithelial compartment containing multiple cell states at the interface with activated fibroblasts (Fig. 6I). We next examined how epithelial-stromal communication changed across the perifocal gradient and TNC^+^ core using ligand–receptor analysis. Epithelial-to-stromal interaction frequency increased towards the core (Fig. 6J-K, S6E-G & Table S4D-E, H-I), with more unique PanCK^+^-to-PanCK^−^ interactions in core-adjacent (R1) regions than in distal (R3) regions (Fig. 6J, S6E & Table S4D). Distal regions favoured inflammatory and chemotactic signals, whereas core-adjacent regions were enriched for matrix-associated and profibrotic pathways. At the centre of this gradient, the TNC^+^ core exhibited features of a signalling hub, with elevated outgoing signalling strength for matrix-associated pathways relative to surrounding regions, whereas inflammatory and immune-associated pathways contributed proportionally less to communication within the core (Fig. 6K, S6F & Table S4E, H).

Within the perifocal epithelium, MIF and APP emerged as dominant outgoing signals directed toward stromal compartments, with MIF-CD74 interactions detected across multiple perifocal regions (Fig. S6F-G & Table S4H-I), demonstrating spatially extended epithelial-stromal communication. Reciprocally, stromal compartments exhibited signalling via non-canonical WNT (WNT5A/WNT5B) and FN1 pathways targeting epithelial receptors (Fig. 6K, S6F-G & Table S4E, H-J), supporting bidirectional epithelial-stromal crosstalk. These signalling interactions occurred alongside loss of epithelial homeostatic regulators and activation of TP63- and SMAD-associated programmes in perifocal regions (Fig. 6C & Table S4B), features of basal-like and matrix-remodelling epithelial states. The dense signalling architecture and matrix-associated interaction networks within these regions suggest that TNC^+^ cores remain tightly coupled to surrounding tissue states rather than representing isolated or biologically inert lesions.

In parallel with increasing epithelial-stromal signalling, fibroblast composition shifted markedly, with activated fibrotic fibroblasts representing the dominant population within the TNC^+^ core (Fig. 6L & Table S4C). The core showed prominent expression of CTHRC1 and CCN2 (Fig. 6B,M-N), together with a high-density interaction network dominated by COL1A2 and FN1 ligands interacting with ITGAV-ITGB8 integrin heterodimers, CD44, and SDC4 receptors on neighbouring cells (Fig. S6G & Table S4I).

These regions also exhibited the highest expression of fibrillar collagens (Fig. 6B,F), together with enrichment of collagen maturation and matrix-stabilisation programmes (Fig. 6G-H; Fig. S7A-B), supporting transition to a dense and structurally stabilised matrix state. Three-dimensional mapping further localised expression of genes involved in collagen biosynthesis and matrix stabilisation to TNC^+^ cores (Fig. 6M-Q; Fig. S7C-H).

Together, these data support an inferred spatial trajectory of human lung fibrogenesis. Increasing alveolar septal density marks coordinated collapse of the alveolar-capillary unit. Perifocal zones exhibit bronchiolisation-associated epithelial remodelling, provisional matrix deposition and altered epithelial-stromal communication, converging on TNC^+^ cores enriched for activated fibroblasts, collagen maturation and matrix stabilisation. Functionally integrated with these perifocal regions, the cores emerge as local signalling hubs where epithelial-stromal crosstalk, fibroblast activation and matrix consolidation intersect, coupling local fibrotic remodelling to broader tissue-state transitions.

### Atlas-defined spatial tissue states capture distinct and clinically relevant dimensions of fibrosis

Finally, to assess the clinical relevance of spatial tissue states identified by our multimodal atlas, we applied atlas-derived gene signatures of alveolar-capillary integrity, perifocal epithelial remodelling and stromal remodelling (Table S5A) to independent IPF cohorts. In an IPF lung tissue cohort (Lung Genomics Research Consortium, n=157)^38^, higher perifocal epithelial and stromal remodelling gene signature scores were significantly associated with reduced pulmonary function (DLCO% predicted and FVC% predicted), whereas the alveolar– capillary integrity signature was associated with preserved pulmonary function (Fig. 6R, Fig. S7I). Across three independent bronchoalveolar lavage cohorts (n = 176)^39^, higher perifocal epithelial and stromal remodelling gene signature scores were significantly associated with mortality after adjustment for disease severity (GAP index), whereas the alveolar–capillary integrity signature showed no relationship with mortality (Fig. 6S, Fig S7J-K). Consistent with these findings, genes within the perifocal epithelial and stromal remodelling signatures were enriched among circulating proteins associated with mortality in an independent IPF plasma proteomic study (n = 1226)^40^ (Fig. 6T, Table S5B-C).

## Discussion

We establish a multimodal 3D framework for human lung fibrosis that embeds molecular programmes within intact microarchitectural context, enabling tissue microarchitecture itself to be interrogated as a biological variable. This approach reveals spatially organised transitions in tissue structure, extracellular-matrix state, and cellular identity that culminate in matrix-stabilised fibrotic microenvironments.

Our findings support a model in which fibrosis progression is governed by progressive loss of the structural and signalling boundary conditions that normally maintain alveolar homeostasis. Using XRH-derived tissue density as a structural metric, we identify collapse of the alveolar-capillary exchange unit as a defining axis of fibrogenic injury in human lung fibrosis. Increasing septal density was tightly associated with coordinated depletion of alveolar type 1 (AT1) epithelial cells, capillary endothelial cells, and basement-membrane components, indicating coupled failure of the epithelial-endothelial scaffold required for gas exchange. The strong correlation between loss of AT1 and capillary endothelial populations further supports the concept that septal collapse reflects failure of an integrated alveolar-capillary unit.^41,42^ Although epithelial injury, endothelial dysfunction, and basement-membrane abnormalities have been implicated in pulmonary fibrosis, these processes have largely been examined independently or in two-dimensions.^42–44^ Our 3D maps directly link their spatially coupled loss to septal microarchitectural collapse, extending morphometric models of alveolar derecruitment in IPF.^45^

Basement-membrane disruption was central to this transition. Spatial transcriptomic analyses and independent extracellular-matrix proteomics showed loss of laminin and collagen IV basement-membrane components as interstitial fibrillar matrix increased, directly linking specialised basement-membrane loss, including COL4A3 and LAMA3, to density and architectural deterioration.^46,47^ High-density alveolar septae did not exhibit enrichment of collagen maturation or matrix stabilisation programmes, suggesting that early architectural failure precedes the emergence of mature fibrotic matrix rather than merely reflecting diffuse collagen accumulation.

These structural changes are likely to have profound consequences for epithelial and endothelial cell fate. Basement membranes are not passive structural scaffolds but active regulators of cellular identity, polarity, and differentiation.^48^ Loss of laminin-rich environments that support AT2-to-AT1 differentiation may therefore destabilise alveolar epithelial identity while simultaneously altering endothelial homeostasis and epithelial-mesenchymal signalling. In this context, collapse of the alveolar-capillary scaffold may represent loss of the physical and biochemical framework required for regenerative alveolar repair.

Regions surrounding areas of active fibrogenesis formed perifocal zones that behaved as transitional repair microenvironments rather than abrupt pathological boundaries. These niches exhibited induction of basaloid/airway-lineage programmes, proteolytic matrix remodelling, altered epithelial-stromal communication, and progressive fibroblast activation. Within perifocal epithelial regions, loss of alveolar epithelial regulatory programmes was accompanied by activation of basal-like and bronchiolisation-associated states, supporting the concept that epithelial reprogramming precedes and surrounds focal matrix accumulation.

This epithelial transition occurred alongside induction of basal keratins, proteolytic programmes including MMP7, and altered epithelial-matrix interactions, indicating that epithelial remodelling develops within an actively changing extracellular environment. Stromal compartments simultaneously exhibited progressive activation of matrix-remodelling programmes characterised by fibrillar collagen deposition and increased matrix-remodelling enzyme expression. Together, these observations suggest that perifocal regions represent spatially organised zones of failed epithelial-mesenchymal coordination in which repair responses remain persistently activated but fail to restore alveolar architecture.

More broadly, our findings suggest that bronchiolisation may represent a stereotyped structural adaptation to collapse of the alveolar microenvironment rather than simply an aberrant lineage state. As the compliant alveolar scaffold deteriorates through combined loss of basement-membrane integrity, epithelial-endothelial coupling, and capillary support, epithelial programmes progressively shift toward mechanically and transcriptionally distinct airway-like states. This transition is accompanied by coordinated activation of basal epithelial regulatory programmes together with altered matrix signalling and adhesion pathways. While such airway-like remodelling may transiently stabilise damaged tissue architecture, it does so at the expense of restoring the thin, tension-balanced alveolar surface required for gas exchange.

Our findings further indicate that these transitional niches converge on spatially restricted TNC^+^ cores that function as matrix-stabilised signalling hubs within the fibrotic trajectory. These regions, corresponding histologically to fibroblast foci, exhibited maximal enrichment of fibrillar collagen synthesis, collagen maturation programmes, and activated fibrotic fibroblast populations. In contrast to perifocal regions characterised by active but imbalanced matrix turnover, TNC^+^ cores demonstrated features of structurally stabilised fibrotic matrix, including enrichment of collagen post-translational modification and cross-linking programmes together with matrix-associated signalling networks centred on CTHRC1^+^ fibroblast populations.

The interaction networks surrounding TNC^+^ cores are consistent with mechanically reinforced fibroblast-matrix signalling that can sustain integrin-mediated TGF-β activation and matrix consolidation.^31,49,50^ Their dense signalling architecture argues against them reflecting inert endpoints; instead, they remain coupled to surrounding stromal, immune and vascular remodelling and may increasingly resist regenerative repair.

How diverse upstream determinants converge on the architectural failure of the alveolar-capillary exchange unit remains to be defined. Genetic and cellular susceptibilities, including MUC5B promoter variants, telomere dysfunction, and surfactant-trafficking abnormalities, may impair the regenerative capacity required to restore alveolar epithelial and basement-membrane integrity following injury.^51,52^ Within this framework, the collapse of the alveolar-capillary scaffold may represent a common structural transition through which heterogeneous pathogenic drivers are channelled into maladaptive fibrogenic repair trajectories.

These findings have potential implications for regenerative intervention strategies. If alveolar-capillary unit loss represents an early organising event in fibrogenesis, approaches aimed at preserving basement-membrane integrity, epithelial-endothelial coupling, or laminin-rich alveolar microenvironments may help maintain regenerative alveolar repair states. Conversely, matrix-stabilised fibrotic cores may represent less reversible tissue states reinforced by mechanotransductive and integrin-mediated signalling networks. These observations raise the possibility that therapeutic responsiveness may differ across distinct spatial stages of fibrotic remodelling. The distinct clinical associations of these spatial tissue states further support this model. Perifocal epithelial and stromal remodelling signatures were associated with impaired lung function and mortality across independent IPF cohorts, whereas loss of alveolar-capillary integrity was associated with physiological impairment but not mortality, suggesting that structural collapse and maladaptive repair capture complementary dimensions of disease.

Beyond the alveolar compartment, our atlas demonstrates that vascular remodelling forms an additional layer of spatial organisation within fibrotic tissue. Fibrotic vascular compartments exhibited loss of endothelial and smooth-muscle homeostatic programmes together with increased biosynthetic activity and altered stress-response signalling. Moreover, transcriptional programmes linked to proteostasis, inflammatory signalling, and cellular stress responses varied according to three-dimensional proximity to remodelled vascular structures. These observations suggest that vascular remodelling and surrounding fibrotic niches become spatially coupled following loss of alveolar-capillary integrity. The emergence of venous and lymphatic endothelial populations within perifocal regions further supports the concept that endothelial identity itself becomes spatially re-patterned within remodelled tissue microenvironments.

This study has several limitations. The technically and resource-intensive multimodal workflow was applied to a limited number of donors. However, extensive within-lung sampling enabled detailed spatial reconstruction, and the findings were supported by multiple independent cohorts and orthogonal datasets. The inferred trajectory recapitulated independently derived gene-expression programmes, extracellular matrix transcriptional changes were concordant with protein dysregulation in independent decellularised-lung proteomic datasets, and atlas-derived spatial signature analyses demonstrated that these tissue states capture distinct and clinically relevant dimensions of fibrosis across independent patient cohorts. Larger disease-stratified cohorts will be required to study later disease stages and fibrotic subtypes. Cross-sectional sampling also cannot establish temporal progression or causality. Experimental perturbation will be required to determine whether alveolar-capillary collapse drives remodelling and whether transitional repair states remain reversible before matrix stabilisation.

Taken together, our findings support a model in which collapse of the alveolar-capillary exchange unit represents loss of the regenerative structural boundary conditions required to maintain alveolar homeostasis, redirecting tissue repair toward spatially organised maladaptive epithelial, stromal, vascular, and immune states. These spatially organised transitional niches progressively converge on stabilised fibrotic cores reinforced by extracellular-matrix consolidation and altered tissue architecture. By anchoring molecular programmes within intact 3D microstructure, this atlas provides a framework for understanding fibrosis as an emergent spatial tissue-state process rather than solely a consequence of isolated molecular pathway activation.

## Acknowledgments

This project was supported by the Medical Research Council (MR/Y002989/1 and 2747520), Boehringer Ingelheim, NC3Rs (NC/V002384/1), the Wellcome Trust (100638/Z/12/Z), and the AAIR Charity. J.B. acknowledges the support of the NIHR Southampton Biomedical Research Centre. The project design schematic in Figure 1A was created with BioRender.com. We thank Carine Fixmer, Maria Lane, Benjamin Johnson, and the nurses of the Southampton Biomedical Research Unit for their help in the collection of human samples, supported by the Wessex Clinical Research Network and the National Institute for Health Research, UK. The authors acknowledge the Faculty of Medicine BIO-R: Bioinformatics Research Facility at the University of Southampton for their support and assistance in this work. The XRH facility used in this research was launched by the Wellcome trust ‘Foundations for routine 3D X-ray histology’, grant 212940/Z/18/Z. XRH data processing was supported by a grant from NVIDIA and utilized an NVIDIA RTX A6000. This research used equipment purchased by the National Research Facility for Lab X-ray CT (NXCT) through EPSRC grant EP/T02593X/1. Research in K.N. lab is supported by funding from Asthma, Allergy and Inflammation Research Charity, DatlaLab and BioFAIR. This research was funded in whole or in part by the Wellcome Trust (209553/Z/17/Z and 227835/Z/23/Z). J Jacob was also supported by the NIHR UCLH Biomedical Research Centre. This project has also been made possible in part by grant number CZIF2024-009938 from the Chan Zuckerberg Initiative Foundation and the Rosetrees Trust (CF-2023-M-2\105).

## Author Contributions

M.G.J. and D.E.D., conceived the study. J.B., M.T., K.G., P.S., S.R., O.K., K.N., Y.W., R.P., D.E.D., and M.E. supported study design and interpretation. S.K.R., P.J.B and O.L.K performed XRH assays. R.P. and M.E. performed GeoMx assays and analysis. G.H. and G.M. supported GeoMx studies, S.J. and J.N. performed CosMx assays. S.K.R. and J.B., performed 3D data integration. A.S. and J.J. supported airway and blood vessel segmentations. J.B., G.A. and K.N. performed all integrative bioinformatic analyses. I. R., J.T., A.F.V., and V.B. supported bioinformatic analysis. J.P., R.A., S.M., D.B., D.J., and L.S. supported multi-modal analyses. A.A., S.J.C, C.H.O, B.G.M, T.W. supported tissue collection. J.N. and J.W. supported tissue processing. A.F., G.T., S.J. supported pathology review and interpretation. J.B., S.R, D.E.D, K.N. and M.G.J drafted the manuscript. All authors read and approved the manuscript.

## Declaration of generative AI and AI-assisted technologies in the writing process

During the preparation of this manuscript, Microsoft Copilot was used to assist with proofreading and editing for readability. The authors reviewed and edited the output as necessary and take full responsibility for the content of the published article.

## METHODS

### Data and code availability

Data can be rendered in MoBIE, a Fiji plugin for multi-modal big image data sharing and exploration,^27,28^ as interactive 3D atlases containing spatially registered volumes of interest, allowing exploration of XRH data combined with 3D spatial RNA and protein abundance visualisation (Fig. S3). The processed multimodal data including the interactive 3D MoBIE atlas are available at doi/10.5281/zenodo.20763804. The raw and processed data are uploaded to GEO (GSE345143 and GSE346080). All original code, supplementary and associated data is available at GitHub (https://github.com/Natarajanlab/3D-IPF-Atlas).

This paper also analyzes existing, publicly available data. The spatial transcriptomic laser-capture microdissection dataset of IPF and control lung tissue is available via NCBI Gene Expression Omnibus under accession GSE169500;^9^ the gene expression profiling of chronic lung disease for the Lung Genomics Research Consortium under GSE47460;^38^ the BAL cell gene expression profiles in IPF under GSE70866.^39^ The digital spatial profiling dataset is available within the supplementary materials of Eyres et al., 2022, *Cell Reports* 40:111230 (https://doi.org/10.1016/j.celrep.2022.111230).^4^ The circulating proteomic dataset is available within the supplementary materials of Oldham et al., 2024, *Am. J. Respir. Crit. Care Med* 209: 1111-1120 (https://doi.org/10.1164/rccm.202301-0117OC).^40^

### Study participant details

Clinically indicated diagnostic surgical lung-biopsy specimens from patients who subsequently received a multidisciplinary diagnosis of IPF according to international consensus guidelines.^53^ Specimens had been diagnosed as showing a typical usual interstitial pneumonia pattern by at least 2 expert lung histopathologists, including areas of morphologically preserved lung tissue including alveolar septae, areas of active fibrogenesis (fibroblastic foci) as well as established fibrosis. Control lung tissue was from macroscopically normal lung sampled remote from the cancer site in age and sex-matched patients undergoing lobectomy surgery for early-stage lung cancer. Details of donor characteristics (n=3 per group) including age, sex, and disease status are provided in Table S6A. Formalin fixed paraffin embedded (FFPE) tissue blocks had received standard processing with fixation in neutral buffered formalin for 48 h and embedding in paraffin wax and tissue blocks had been archived for less than seven years. The study was approved by the Southampton and South West Hampshire and the Mid and South Buckinghamshire Local Research Ethics Committees (ref. 07/H0607/73, 14/SC/0186), and all participants gave written informed consent.

### 3D X-Ray Histology Scanning

FFPE tissue samples were scanned non-destructively using a customised micro-computed tomography scanner optimised for 3D X-ray histology based on Nikon’s XTH225ST system (Nikon Metrology, Tring, UK).^15,20^ The scanner was operated at 60 kVp/115 μA (6.9 W), using a molybdenum (Mo) target without beam pre-filtration. The voxel size ranged from 4.9µm to 8.7μm to allow for a larger field of view for the larger samples. Imaging parameters were as follows: 3501 projections were taken over the 360° rotation, with four frames per projection averaged to improve the signal-to-noise ratio. The exposure time for each projection was 1415 ms and the detector’s gain was set to 24 dB. After CT acquisition, the data were reconstructed into 32-bit volume files by means of Nikon’s own reconstruction software (CT Agent, Nikon Metrology, Tring, UK), which used a filtered-back projection algorithm. The reconstructed 32-bit raw image volumes were then converted to 16-bit within the Fiji distribution of ImageJ,v2.16.0/1.54p for visualisation.^28^

Each tissue block was scanned using the same XRH acquisition and reconstruction protocol and system to minimise variability, with magnification adjusted to capture each tissue block in full. Reconstructed greyscale values were normalised across datasets to enable comparison between samples. Normalised XRH-derived relative tissue density (tissue mass density) refers to a relative attenuation-based metric from the reconstructed volumes, reflecting local differences in tissue structure. The same processing pipeline was applied to all control and IPF samples. Paraffin intensity distributions were comparable across tissue blocks, whereas tissue intensity distributions retained the expected biological variation.

### Spatial analysis of TNC and ACTA2 expression

To investigate TNC and ACTA2 spatial expression, we re-analysed two complementary spatial transcriptomic datasets of IPF and control lung tissue that we previously generated. The first was a laser-capture microdissection (LCM) RNA-sequencing dataset (GSE169500) in which FFPE lung tissue was microdissected from three region types - alveolar septae from non-fibrotic control lung donors (n = 10), fibroblast foci from usual interstitial pneumonia/IPF lung donors (n = 10), and adjacent non-affected alveolar septae from the same IPF lung tissue (n = 10) - followed by total RNA extraction, cDNA library preparation using the Ion AmpliSeq Transcriptome Human Gene Expression kit (Life Technologies, Paisley, UK), and sequencing on an Ion Torrent Proton platform.^9^ Reads were aligned to human reference genome GRCh38 using Kallisto and counts per million mapped reads (CPM) were calculated. The second was a digital spatial profiling (DSP) dataset generated using the NanoString GeoMx Cancer Transcriptome Atlas on FFPE IPF and control lung tissue, with morphologically defined regions of interest and and transcript counts quantified within each ROI.^4^ The LCM dataset is available via NCBI GEO under accession GSE169500; the DSP dataset is available within the supplementary materials of Eyres et al.^4^

### Spatial profiling (Bruker GeoMx)

#### Spatial transcriptomic profiling

Tissue sections were processed for digital spatial profiling on the Bruker GeoMx platform. IPF & control blocks were processed pairwise. FFPE blocks were sectioned at 5µm depth and mounted on poly-L-lysine coated glass slides. Sections were inspected for distortion and replaced with an adjacent section if substantial sectioning artefacts were identified.

Ten sections (≈100µm between each section, total sectioning depth ~1mm) per block, were processed for spatial transcriptomic profiling. In brief, sections were probed with the human Whole Transcriptome Atlas (WTA) panel (Bruker, GMX-RNA-NGS-HuWTA-4). Tissues were stained for cytokeratin (Bruker, GMX-RNA-MORPH-HST-12), CD45 (Bruker, GMX-RNA-MORPH-HST-12) and Tenascin C (Abcam, ab271877) to identify epithelial cells, immune cells and areas of active fibrogenesis respectively. ROIs were selected and segmented as described below. Photocleaved oligonucleotide barcodes were collected from a total of 1,450 ROIs. Oligonucleotides were quantitated via Illumina next generation sequencing. Libraries were sequenced using an Illumina NextSeq 1000 (P2 Reagents (100 cycles), Illumina, #20046811).

#### Spatial protein profiling

A slide adjacent to that used for spatial transcriptomics was processed for spatial protein profiling using the Bruker GeoMx platform. Within 2 weeks of microtomy, sections were probed with a 77-plex protein panel consisting of human: Immune Cell Profiling Panel (#121300101), IO Drug Target Panel (#121300102), Immune Activation Panel (#121300103), Immune Cell Typing Panel (#121300104), MAPK signalling panel, Cell Death Panel (#121300112), PI3K/AKT Signalling Panel (#121300113). Sections were concurrently stained with the same morphology markers used for spatial transcriptomic profiling to guide ROI selection and segmentation.

### ROI selection

Airways, blood vessels and alveolar septae were sampled from both control and IPF sections, while immune infiltrates and areas of active fibrogenesis (TNC^+^ cores and their perifocal regions) were unique to IPF sections. ROI definitions are given below. Where compartmental segmentation was required, marker-positive and marker-negative AOIs were generated using a common mask-based workflow. Fluorescence-intensity thresholds were applied to the relevant marker channel to generate binary masks.

The following ROI types were selected, as applicable:

#### Alveolar septae

In IPF and control tissues, square ROIs with 450µm side-length were selected in regions with typical alveolar septae morphology.

#### Airways

For each IPF and control block, airways belonging to bronchovascular bundles spanning the depth of tissue samples were identified by reviewing the XRH volume. ROIs were drawn manually around the epithelial tissue and was segmented into panCK^+^ and panCK^+^ compartments according to panCK staining.

#### Blood vessels

In each IPF and control tissue block, arterial vessels adjacent to airway ROIs were selected. ROIs which bisected the width of vessel walls were manually drawn to enrich vascular intima (BVi) and adventitia (BVa) layers.

#### TNC^+^ Cores

ROIs measuring 660 × 785 µm were centred on TNC^+^ cores identified by visual inspection of the immunofluorescence images. Each ROI was segmented into areas of interest (AOIs) by first creating a TNC^+^ mask, defined as pixels with staining intensity above an image-specific threshold. The mask was subsequently smoothed and subjected to hole filling. Three 90 µm-wide concentric region masks were created by sequential outward expansion from the TNC^+^ mask, producing Region 1 (adjacent to the TNC^+^ mask), Region 2 (intermediate), and Region 3 (most distal from the TNC^+^ mask). Each concentric region was further segmented using a pan-cytokeratin (panCK) staining intensity threshold to generate panCK^+^ and panCK^−^ binary masks as with the TNC^+^ mask above, enriching for epithelial and non-epithelial areas, respectively. All intensity thresholds were generated with reference to a single reference image. After digital spatial profiling, tissue sections were stained using haematoxylin 7211 (Fisher Scientific, #10034813) and counterstained with eosin Y (Fisher Scientific, #12677756). Sections were mounted in ProLong Gold Antifade mountant (Thermo Fisher Scientific), then imaged using a Zeiss Axio Scan Z1. After the DSP protocol was complete each section was examined and ROI annotation confirmed by a lung histopathologist (A.F.)

#### Immune infiltrate

Circular ROIs 500µm in diameter were selected in regions enriched for **CD45^+/-^**immune infiltrates. ROIs were segmented into CD45^+^ and CD45^+^ compartments according to CD45 staining, using a binary mask-based workflow identical to that used for panCK and TNC staining segmentation, selecting separate compartments based on whether they were above or below an image-specific threshold. Immune infiltrates closest to sampled TNC^+^ cores were selected.

### Data processing and QC

FASTQ files were processed using the GeoMx NGS Pipeline (Bruker, MAN-10118-03) before undergoing QC for minimum raw reads (1000), sequencing saturation (>50%) and percent aligned, stitched, and trimmed reads (>80%). ROIs with less than 10% of genes expressed were removed. QC processing was performed using the GeomxTools R package.^54^ Following QC, 193 ROIs were excluded, leaving a total of 1,257 ROIs. Batch correction was applied using the ComBatSeq R package.^55^ Following batch correction, Q3 normalisation was applied to count data. Normalized counts were exported and processed further using R. QC and normalisation of spatial protein data were performed via Bruker-DSP Analysis Suite (Bruker, v3.1.0.222). Probes with low signal-to-background ratios (NF1 (not present in probe lot), CD80 & GITR) were excluded from analysis. Raw probe count data was then normalised to negative IgG counts for downstream analysis. Following QC, 1,362 protein ROIs were included.

Dimensionality reduction of normalised GeoMx data was performed using t-distributed stochastic neighbour embedding (t-SNE).

### Cell profile matrix and spatial deconvolution

A cell profile matrix was generated from a published lung spatial RNA-sequencing dataset as a reference for cell type signatures.^14^ Mean expression of all genes for each defined cell type was calculated. This set of marker genes was then used as the cell profile matrix for deconvolution. Spatial deconvolution was performed using the SpatialDecon R package.^56^ The spatialdecon() function was used to deconvolute the bulk gene expression data from individual regions of interest into the proportions of different cell types associated with each ROI. Background values were derived from the negative probe expression values.

### CosMx Spatial Molecular Imager Assays

*In situ* image-based quantification of ~6000 genes (CosMx™ Human 6K Discovery Panel, #121500041) were performed according to manufacturer’s instructions using the CosMx Spatial Molecular Imager, with morphology IF markers CD45, CD68, panCK, CD298/B2m, DAPI (CosMx™ Human Universal Cell Segmentation Kit (#121500020) & CD68 marker kit (#121500022). The entire available tissue for each section was imaged through sequential field of view acquisition.

Following image acquisition, cell boundaries were determined using the AtomX cell segmentation algorithm, using configuration A (non-neuro human tissue). Data were processed according to the default AtomX pipeline. CosMx flat files were downloaded from AtomX, and processed using the Seurat R package. Per-cell gene expression was log normalised, and dimensional reduction was performed using Uniform Manifold Approximation Projection (UMAP) plotting. Cells were annotated using the InSituType algorithm with cell-type signatures derived from the same Vannan et al. lung spatial transcriptomic reference used for SpatialDecon analysis above.^14^ Spatial polygon maps of cell types and gene set variation analysis (GSVA) scores were generated in R using polygon flat files.

### XRH & Spatial Profiling Data Integration

The XRH volume was used as the reference for integrating transcriptomic and protein expression data, with ROIs digitally labelled within the XRH volume, so enabling establishment of a 3D multi-modal resource. The methods applied for this integration are described below. The intact XRH volume served as the geometric reference because histological sectioning can distort tissue geometry. Analyses were performed at ROI/VOI and compartment level rather than voxel by voxel, thereby limiting the effect of small residual local misalignments caused by section stretching, compression or deformation.

### XRH histology alignment

In Fiji, 16-bit XRH volumes were rotated and resliced using bicubic interpolation so that the x-y plane approximately matched the sectioned face of each block. The volume was then rigidly resliced to align precisely with the first stained DSP section using the three-point alignment tool in the Slice module of Avizo 3D (v. 2024.1) (Thermo Fisher Scientific).^57^

### Segmentation of DSP Regions of Interest (ROIs) into XRH data

Avizo 3D was used to generate ROIs within each XRH volume. One project was used per tissue block, a single label file was used for each equivalent DSP ROI set, and a separate material was given for each ROI. Each ROI material was created in the same order within each label file to allow data visualisation automation. For example, for control samples, the material list was assigned Alveolar ROIS (1-4), AW (5-6) and then BV (7-8). A pseudo-ROI was placed within the file if an ROI was missing from the DSP slide. Due to the precise alveolar pattern registration between the XRH data and the DSP-stained slides, the ROIs could be manually segmented accurately.

Following the approach of the DSP ROI selection, for the control samples the alveolar septae ROIs were segmented with a square brush tool. Blood vessels ROIs (BV) were segmented to include the whole cross-section of the blood vessel, and airways.

Following the approach of the DSP ROI selection for the IPF samples, the AS, BV and AW ROIs were segmented in the same way as the control tissue. The Immune infiltrate ROIs were segmented with a circular brush tool. For segmentation of areas of active fibrogenesis, the central TNC^+^ core and the three concentric perifocal regions were all segmented as separate materials as follows. Each TNC^+^ core was manually segmented into the XRH in the same pattern defined by the DSP-stained section and added to the core material. The first perifocal region was defined by dilating the outer circumference of the TNC^+^ core region by 90µm. The second and third regions were generated similarly but dilated from the previously generated region’s boundary. The outer box restriction to the Core ROIs was not adhered to during the ROI transfer. This resulted in some outer regions being complete, whereas within the DSP ROIs, they were disjointed. This was necessary to allow for 3D volume of interest generation. Where any ROI boundary overlapped, the following order of priority was given: active fibrogenesis, Immune, Airways, Blood vessels, and Alveolar Septae. The lower priority ROI had material subtracted from it.

### Volume of Interest (VOI) generation

All ROI to VOI generation was made within Avizo software unless otherwise stated. All the ROIs were extended up and down the Z-axis, keeping the DSP ROI the central slice of the VOI until the total Z-axis dimension equalled 100µm. Where any extension intersected with another VOI in the Z axis, volume was removed equally from each intersecting VOI. The BV and AW ROIs were extended up and down the Z-axis using their respective structures within the XRH volume. This was achieved through manual segmentation and Avizo’s interpolation algorithms. The alveolar septae and immune ROIs did not have any apparent reference point within the XRH volume to dictate the extension direction; therefore, the same ROI shape in the X-Y plane was extended in the z-axis.

From these 3D files, 100 µm VOIs were extracted and placed back into the original label file to make the 3D VOIs for each 2D core identified from the DSP methods.

The label files representing the DSP analysed slides, now containing 3D VOIs, were exported from Avizo 3D in a 3D raw format. In Z-axis order, each label file was imported into Fiji as an 8-bit image file and added to the 3D VOI manager within the 3D suite plugin.^58^ Once every VOI, within all the label files, is placed within the 3D VOI manager, an index number, volume and X,Y and Z coordinates of every VOI can be generated within each tissue sample.

An ImageJ macro-language script was written that automatically assigns voxels within each 3D manager VOI selection with an 8-bit intensity value based on a lookup table derived from the GeoMx gene and protein expression data. The script used for voxel assignment and visualisation is available in the project GitHub repository (see Code Availability). This was applied to an empty image space of the same dimension as the XRH image volume. Using a linear index value of 1-220 for IPF tissues and 1-80 for the control samples (1-64 for Cnt002), a VOI-indexed image was generated for each tissue.

Expression data was scaled to a range of 1-255 (1= no expression, 255 = high expression) relative to all other genes/proteins across the entire dataset. This was to allow control and IPF visualisation simultaneously, and the data to be relative to each other. The value of zero was used for VOIs that did not return any data. Individual .csv files were generated for each sample to include combinations of sub-ROI cell segmentation e.g. PanCK^+/-^, CD45^+/-^and for the blood vessel’s outer (adventitia)/ inner (intima).

### Visualisation of Omics data within VOIs

For each gene or protein, the VOI-indexed image, ImageJ VOI-intensity editing script and corresponding omics CSV file were combined to generate an 8-bit spatial abundance volume in Fiji.

Eight-bit RNA expression volumes for *TNC, PTPRC* and *ACTA2* were generated as test datasets because their spatial patterns were expected to be readily interpretable. Volumes were exported to Avizo and displayed using the Ironbow colour palette to facilitate comparison across expression intensities and improve accessibility for viewers with colour-vision deficiency.

### 3D structural Segmentation

Airway and vascular segmentation were achieved through a combination of an artificial intelligence framework for segmenting airway and vascular structures from lung XRH images as well as manual segmentation techniques. The framework uses a set of nnUNets^59^ as a backbone and was trained using manually annotated lung structures at Hierarchical Phase Contrast Tomography (HiP-CT) and XRH images by radiologists. The XRH images were pre-processed before the inference by removing the outliers by scaling their intensities. The automated method’s result segmentations were manually revised and corrected. Airway structures were removed due to some inaccuracies in this segmentation in the current development of the model. Other vascular structures were dilated and then eroded within Avizo by 2 voxels. Any label object data that exceeded 5000 voxels were retained; all other structures were deleted. Any labels that exceeded 5000 voxels that were not vascular structures were deleted. Manual segmentation was undertaken on all other visible vascular structures down to the spatial resolution limit of the XRH datasets.

Airways were segmented using manual techniques. Anatomically normal airways were segmented to the terminal bronchioles that retained a lumen. Limited airway data was segmented across the samples due to their peripheral nature and the lack of normal airway structures in the IPF samples. Therefore, the time commitments of manual segmentation for these structures were justified for the accuracy it gives.

Tissue structures were segmented using automated techniques within Fiji. Thresholding, outlier removal, erosion/dilation tools and the “Keep-largest” function within the MorpholibJ plugin were used.

Whole lung volume segmentation was achieved using the binary tissue segmentation data as a base. Then, alveoli and other inclusions within the tissue were filled in using the 3D suite’s “fill holes” function and MorpholibJ 3D morphological functions 3D erode/dilate and 3D median filters.

### Structural analysis of VOIs using XRH data

To confirm accuracy of segmentation, all structures were reviewed by an expert histopathologist (AF) and the finalised segmentations proceeded to downstream analysis.

The VOI-indexed image for each sample was loaded in Fiji then added to the 3D ROI manager within the 3D suite plugin. The 16-bit XRH volume of the corresponding sample was loaded into Fiji.

All VOIs within the 3D manager were then selected, and the centroid, volume, and mean greyscale metrics were measured upon the 16-bit greyscale intensity XRH volume giving the X, Y, Z coordinates, VOI Volume and greyscale intensity of XRH data for each VOI analysed. Alveolar greyscale intensity was included in this analysis (if within the VOI extent) to reduce the potential errors acquired in the segmentation of the tissue structure.

Using the 3D local thickness function within the BoneJ plugin^60^ within Fiji, a local thickness map was generated for each tissue volume using the binary segmented tissue volume file as an input. This replaced the XRH volume file as the input for the 3D ROI manager analysis, thus the mean greyscale value measure returned the result of the 3D local thickness map.

3D geodesic distance maps from both the blood vessel and airway segmentation were generated within the MorpholibJ plugin within Fiji.^61^ The input image was the binary vessel or airway image file, respectively, and the mask was the binary tissue volume file. This also replaced the XRH volume file as the input for the 3D ROI manager analysis, thus the mean greyscale value measure returned the result of the 3D geodesic distance maps.

The results of the 3D local thickness maps and geodesic distance maps were 32-bit files thus all background data was assigned N/A, thus not included in the 3D suite analysis.

To acquire the tissue volume fraction for each VOI, the binary segmented tissue volume file was loaded into Fiji and multiplied by the VOI-indexed image. This resulted in an image with only the tissue volume in each VOI-indexed file. The tissue volume for each VOI was, therefore, acquired in the same method as stated above. This divided by the total VOI volume gave a tissue volume fraction for each VOI analysed.

## QUANTIFICATION AND STATISTICAL ANALYSIS

### Statistical analysis and visualisation

Analysis-specific statistical procedures are described in the subsections below and in the corresponding figure legends. Unless stated otherwise, tests were two-sided. Associations between continuous variables were assessed using Pearson’s product-moment or Spearman’s rank correlation, as specified for each analysis. P ^62–64^ values were adjusted for multiple comparisons using the Benjamini-Hochberg false-discovery-rate method where applicable. Graphs were generated using ggplot2 and heatmaps using the pheatmap R package.^65^

### GeoMx differential-expression analysis

Differential gene expression in GeoMx data was assessed using linear mixed-effects models implemented in lme4, with anatomical region as a fixed effect and slide ID as a random intercept.^63^ Regional marker genes were identified using Limma.^64^ Associations of gene expression, cell-deconvolution scores and GSVA scores with XRH-derived tissue mass density were evaluated using cor.test() in R.

### Gene Set Variation Analysis

Gene set variation analysis (GSVA was performed in R using the gsva package and Biological process gene ontology terms, excluding terms with <5 or >200 genes.^66–68^ Endothelial-subtype gene sets were constructed from marker genes reported in the integrated single-cell atlas of human lung endothelial cells by Schupp et al.^69^ Basement-membrane gene sets were obtained from the BMbase resource described by Jayadev et al.^70^ For CosMx analyses, single-cell wound-response matrix scores were calculated from the log-normalised expression matrix. GSVA scores were used for comparisons between anatomical compartments and for correlation with XRH-derived tissue mass density.

### Gene Regulatory Network (GRN) Inference

The pySCENIC pipeline (Python implementation of SCENIC) was used to infer gene regulatory networks with modifications,^71,72^ independently across eight distinct tissue compartments: airway IPF, alveolar IPF, blood vessel IPF, immune infiltrate IPF, TNC^+^ core (comprising TNC^+^ core, PanCK^+^, and PanCK^−^ regions R1-R3), airway control, alveolar control, and blood vessel control.

For each compartment, co-expression modules between transcription factors (TFs) and candidate target genes were first identified using GRNBoost2 (pyscenic grn) with the human TF reference list (allTFs_hg38.txt) ^73^. Modules and indirect targets were subsequently pruned by motif enrichment using pyscenic ctx against two cis-regulatory databases: a 500 bp upstream promoter ranking and a 10 kb TSS-centred ranking (hg19, 7-species conservation, mc9nr) with HGNC motif annotations (motifs-v10nr_clust-nr, threshold m0.001).

Regulon activity per region-of-interest (ROI) was quantified as Regulon Activity Scores (RAS) (pyscenic aucell), yielding a regulon x ROI activity matrix for each compartment. Regulons were filtered to retain those detected in ≥10 ROIs per subanatomical region (>2% of total ROIs) and exhibiting non-zero scores in ≥50% of ROIs. Resulting RAS matrices were used for downstream heatmap visualisation, differential regulon activity comparisons between IPF and control, and identification of compartment-specific TF programmes (Table S6B-C).

### Tissue Density Correlation Analysis

To assess the impact of fibrotic remodelling on regulatory states within alveolar septae, Regulon activity scores (RAS) were independently correlated with scaled tissue mass density for IPF and control ROIs using Spearman’s rank correlation. A pseudoactivity of 1×10^−8^ was added as the RAS for regulons absent in each condition, following union-based harmonisation of regulon sets identified across IPF and control compartments.

For each regulon, a differential Pearson correlation statistic (Δr = r (IPF) − r (Control)) was derived to quantify the shift in density-association between disease and control states. Statistical significance was evaluated using Fisher’s method, with all resulting p-values adjusted for multiple comparisons using the Benjamini-Hochberg (FDR) method. Results were visualised as volcano plots with Δr on the x-axis, −log10(Fisher combined adjusted p-value) on the y-axis, and regulons colored by differential RAS activity on a diverging colormap.

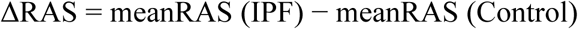

Global differential regulon activity was also computed via two-sided Mann-Whitney U tests. Differential and tissue-correlated regulon activity between IPF and control alveolar regions was visualised using heatmaps stratified by median-split tissue density tiers (Low vs. High; normalised cutoff 0.206), with an additional column representing the differential density correlation (Δr) for each regulon.

Transcriptomic and regulatory profiles were further visualised by summarisation via group-level pseudobulk aggregation (TNC^+^ core) or single-ROI matrices (CD45^+/-^ infiltrates, airways, and blood vessels). For blood vessels and airways, top regulons were visualised using dumbbell plots comparing maximum motif normalised enrichment scores (NES) and TF-target importance shifts across conditions.

### Microenvironment Deconvolution

To estimate cellular composition of spatial ROIs, cell-type enrichment analysis was performed using the Univariate Linear Model (ULM) algorithm implemented in decoupler.^74^ A human lung-specific marker gene reference curated from PanglaoDB was used, restricting inference to cell types with a minimum of three detected marker genes in the dataset. The resulting ULM enrichment scores were aggregated by spatial subregion, z-scored across regions to highlight niche-specific localisation patterns and visualised as heatmaps with statistical significance denoted based on median adjusted p-value per subregion.

### Gene Set Enrichment Analysis and Basement Membrane Dynamics

To investigate extracellular matrix signatures, curated gene sets were evaluated, including Alveolar-Capillary Integrity, Endothelial Stability, and Wound-Response Matrix (Table S6D).^70,75–77^

Pathway enrichment was tested using gene set enrichment analysis (GSEA) implemented in blitzgsea.^78^ Genes were ranked using Welch’s t-test statistics derived from targeted comparisons, enabling both one-versus-rest contextualisation of spatial niches and pairwise comparisons of matched structural regions (IPF vs. control) and intra-IPF focal transition zones (e.g., TNC^+^ core vs. panCK^+/−^ perifocal regions).

### Projection of Alveolar Remodelling Gene Programs onto a Spatial Trajectory

To independently assess perturbed alveolar gene programs across the early IPF microenvironment, we projected the pseudo time-associated gene programs defined by Vannan et al. onto our atlas.^14^ The three gene sets were mapped onto our atlas resulting in Homeostasis (76 shared of 89 genes), Intermediate (30 shared of 34 genes) and Late remodelling (93 shared of 99 genes) respectively. We scored each gene within the gene-set per ROI and plot the mean log-normalized expression. To arrange ROIs across trajectory we first ordered subcompartments from Control, IPF alveolar septae, panCK^−^ perifocal regions (R3, R2, R1), TNC^+^ core, panCK^+^ perifocal regions (R1, R2, R3), followed by scaled tissue mass density. The z-scored mean expression for each ordered ROIs is plotted in Fig. 6I.

To plot trajectory for each gene set, we used generalized additive model (GAM) consisting of natural cubic-regression spline of trajectory position and donor as a fixed-effect covariate, which is fit by ordinary least squares (OLS). The population-level mean trend is obtained by first computing per-donor trends over observed range and averaging model predictions across donors. Data pre-processing, analysis and spline fits were performed in Python using *scanpy* and *statsmodels* libraries.^79,80^

### Clinical cohort analyses of atlas-derived gene expression signatures

Atlas-derived gene signatures were evaluated in independent, publicly available IPF cohort datasets. Compartment-specific remodelling signatures (Table S5A) were defined from the perifocal ring analysis (Fig. 4; Table S4A): an epithelial remodelling signature comprised the 100 most significant genes up-regulated in ring 1 (core-adjacent) versus ring 3 (distal perifocal) within the panCK^+^ (epithelial) compartment. The stromal remodelling signature comprised the corresponding top-100 genes in the panCK^−^ (mesenchymal) compartment, ranked by false-discovery rate. The alveolar–capillary integrity signature comprised canonical alveolar type-1, capillary-endothelial and alveolar basement-membrane identity genes that significantly declined with increasing tissue density.

Signature activity was quantified per sample using singScore, a rank-based single-sample method, applied to expression matrices collapsed from probes to gene symbols (highest-mean probe per symbol)^81,82^. Physiological associations were assessed in lung-tissue transcriptomes from the Lung Genomics Research Consortium (LGRC; GSE47460)^38^. Signature scores were correlated (Spearman) with diffusing capacity (DLCO, % predicted) and forced vital capacity (FVC, % predicted), restricted to IPF/UIP samples (DLCO, n = 145; FVC, n = 157). Prognostic association was assessed in bronchoalveolar-lavage transcriptomes from three prospective IPF cohorts^39^ (GSE70866; Freiburg n = 62, Leuven n = 64, Siena n = 50; 176 patients). Scores were modelled using Cox proportional-hazards regression to estimate the hazard ratio (HR) for mortality per standard deviation increase in signature score. Pooled HRs are reported both unadjusted and adjusted for the GAP index (age, sex, FVC and DLCO). Kaplan-Meier curves were stratified by signature tertile and compared using log-rank tests. Concordance with circulating proteins was assessed using an independent IPF plasma proteomic study (Oldham et al., 2024; Olink Explore 3072; n = 1,226)^40^. All proteins quantified in the study (n = 2,877) were used as the background set for enrichment analysis. For each atlas-derived signature, enrichment of proteins encoded by signature genes was tested against two target sets: (i) all proteins associated with transplant-free survival (FDR < 0.05), and (ii) the subset associated with adverse outcome (hazard ratio > 1), using the hypergeometric test.

### Cell-Cell Communication Inference

Cell-cell communication was inferred using CellChat v2.2.0 applied to two analytical subsets^83^. First, CD45^+^ (immune) and CD45^−^ (parenchymal) cell populations were analysed together to characterise cross-compartment signalling. Second, spatially-defined populations comprising the TNC^+^ core and three concentric peri-focal regions of panCK^+^ (epithelial) and panCK^−^ (stromal) cells were jointly analysed to resolve spatially-graded communication patterns. The human CellChat ligand-receptor database (CellChatDB.human) was used throughout.

Ligand-receptor interactions were identified by subsetting the database to expressed genes (subsetData), followed by detection of over-expressed signalling genes and interactions (identifyOverExpressedGenes, identifyOverExpressedInteractions). Communication probabilities were computed using the truncated mean method (trim = 0.1) with population size correction (computeCommunProb, type = “truncatedMean”, population.size = TRUE), and cell-type groups with fewer than 10 cells were excluded (filterCommunication, min.cells = 10). Pathway-level communication probabilities were subsequently aggregated (computeCommunProbPathway, aggregateNet) and network centrality scores - including outdegree, indegree, and information centrality - were computed to identify dominant senders, receivers, mediators, and influencers (netAnalysis_computeCentrality).

Communication patterns were identified using non-negative matrix factorisation (NMF), with the number of outgoing and incoming patterns selected by inspecting Cophenetic and Silhouette metrics. Four outgoing and three incoming patterns were retained for the spatially-defined dataset; two outgoing and two incoming patterns were retained for the CD45 dataset.

### Visualisation and Comparative Analysis of Cell-Cell Communication

Signalling networks were visualised as chord diagrams, circle plots, and signalling role heatmaps using CellChat built-in functions (netVisual_chord_gene, netVisual_aggregate, netAnalysis_signalingRole_heatmap). To quantify directional communication bias, a pathway-level communication probability matrix was constructed by summing probabilities across all ligand-receptor pairs belonging to each pathway for every directed sender-receiver cell-type pair. Total outgoing communication probability was aggregated across directed pairs within each compartment.

Directional asymmetry was quantified as the log2 ratio of summed outgoing probabilities between the two compartments of interest:

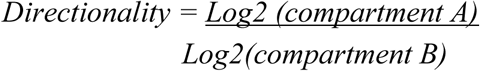

Positive values indicate compartment A dominance; negative values indicate compartment B dominance. A pseudocount of 1×10^−6^ was added prior to log transformation to handle zero values. For the three-compartment spatially-defined analysis, two orthogonal metrics were computed: the log2 ratio of panCK^−^ to panCK^+^ outgoing probability (stromal vs. epithelial dominance), and the log2 ratio of TNC^+^ outgoing probability relative to the mean of the other two compartments (TNC^+^ involvement relative to peri-focal regions), enabling joint visualisation of spatial communication hierarchies.

Pathways were selected through a two-step procedure: an initial candidate set was defined based on computed roles and relevant cell compartments (e.g., canonical immune signalling for CD45^+^ cells; ECM and stromal remodelling pathways for parenchymal and TNC^+^ populations), after which candidates were validated against empirically observed directional scores. Only pathways with total cross-compartment communication probability >0.05 and consistent directional log2 ratios were retained. Pathway-level directional scores were calculated for all inferred signalling pathways. For visualisation, pathways with total cross-compartment communication probability>0.05 were ranked by the absolute magnitude of their directional log₂ ratio. Only representative pathways from the highest-ranking immune, ECM and stromal-remodelling categories were displayed to illustrate compartment-dominant signalling. Pathway-level results (including pathways not displayed) are provided in Table S6E.

