## Supplemental Information for "Three-dimensional mapping of human lung fibrosis identifies spatial coupling between tissue microarchitecture and maladaptive repair states"

#### Supplementary Figures

Figure S1

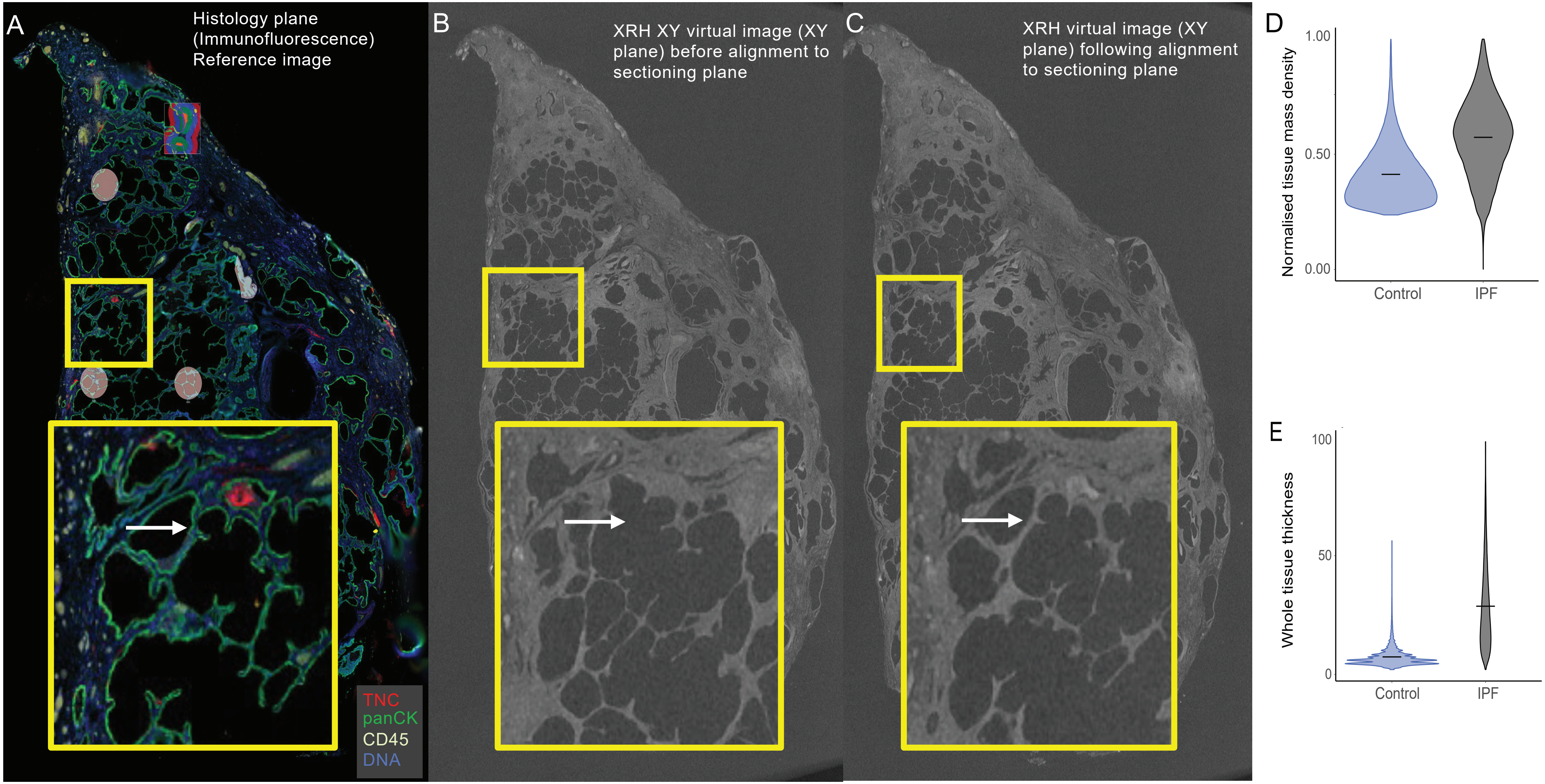
A
Histology plane
(Immunofluorescence)
Reference image
TNC
panCK
CD45
DNA
B
XRH XY virtual image (XY plane) before alignment to sectioning plane
C
XRH virtual image (XY plane) following alignment to sectioning plane
D
Normalised tissue mass density
1.00
0.50
0.00
Control
IPF
E
Whole tissue thickness
100
50
0
Control
IPF

**Supplementary Fig. S1: Co-registration of tissue sections within the 3D X-Ray Histology volume. (A)** Representative immunofluorescence image of an IPF tissue section used for Bruker GeoMx spatial profiling, showing DAPI (nuclei), PanCK (green), TNC (red) and CD45 (yellow) staining. The yellow box outlines the zoomed in area shown below. **(B)** X-Ray Histology virtual image of the same IPF tissue block, showing the XY location of the section in (A) before alignment to the sectioning plane. **(C)** after alignment to the sectioning plane. White arrows in zoomed insets in **(B)** and **(C)** indicate the same anatomical landmark as in (A). (**D**, **E**) XRH-derived structural metrics from control and IPF tissue blocks (n=3 per group): (**D**) voxel-wise tissue mass density and (**E**) whole-tissue local tissue thickness.

Figure S2

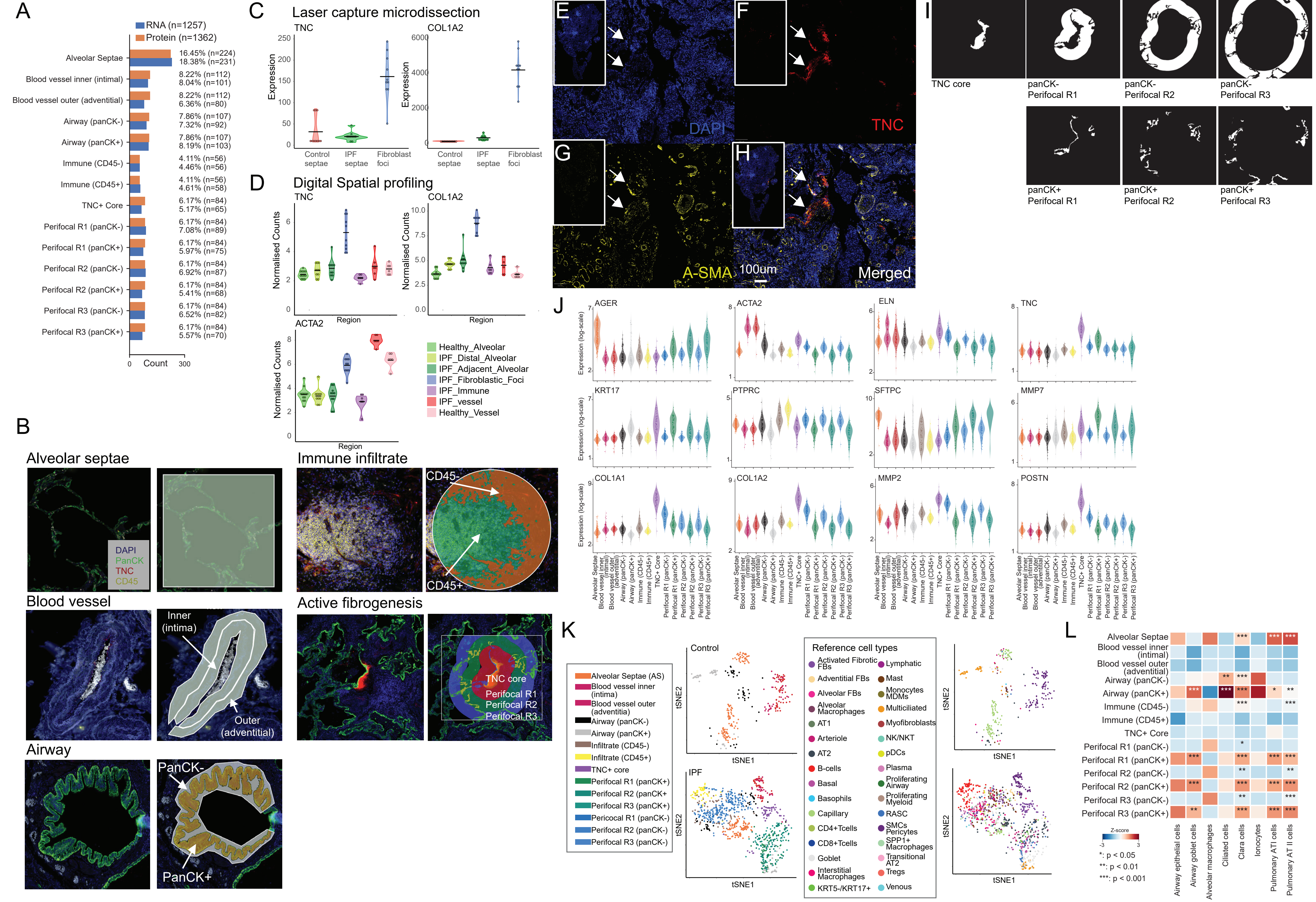
A
RNA (n=1257)
Protein (n=1362)
Alveolar Septae
Blood vessel inner (intimal)
Blood vessel outer (adventitial)
Airway (panCK-)
Airway (panCK+)
Immune (CD45-)
Immune (CD45+)
TNC+ Core
Perifocal R1 (panCK-)
Perifocal R1 (panCK+)
Perifocal R2 (panCK-)
Perifocal R2 (panCK+)
Perifocal R3 (panCK-)
Perifocal R3 (panCK+)
16.45% (n=224)
18.38% (n=231)
8.22% (n=112)
8.04% (n=101)
8.22% (n=112)
6.36% (n=80)
7.86% (n=107)
7.32% (n=92)
7.86% (n=107)
8.19% (n=103)
4.11% (n=56)
4.46% (n=56)
4.11% (n=56)
4.61% (n=58)
6.17% (n=84)
5.17% (n=65)
6.17% (n=84)
7.08% (n=89)
6.17% (n=84)
5.97% (n=75)
6.17% (n=84)
6.92% (n=87)
6.17% (n=84)
5.41% (n=68)
6.17% (n=84)
6.52% (n=82)
6.17% (n=84)
5.57% (n=70)
0
Count
300
C
Laser capture microdissection
TNC
COL1A2
Expression
Control septae
IPF septae
Fibroblast foci
D
Digital Spatial profiling
TNC
COL1A2
ACTA2
Normalised Counts
Region
Healthy_Alveolar
IPF_Distal_Alveolar
IPF_Adjacent_Alveolar
IPF_Fibroblastic_Foci
IPF_Immune
IPF_vessel
Healthy_Vessel
E
F
G
H
DAPI
TNC
A-SMA
100um
Merged
I
TNC core
panCK- Perifocal R1
panCK- Perifocal R2
panCK- Perifocal R3
panCK+ Perifocal R1
panCK+ Perifocal R2
panCK+ Perifocal R3
B
Alveolar septae
DAPI
PanCK
TNC
CD45
Immune infiltrate
CD45-
CD45+
Blood vessel
Inner (intima)
Outer (adventitial)
Active fibrogenesis
TNC core
Perifocal R1
Perifocal R2
Perifocal R3
Airway
PanCK-
PanCK+
J
AGER
ACTA2
ELN
TNC
KRT17
PTPRC
SFTPC
MMP7
COL1A1
COL1A2
MMP2
POSTN
Expression (log-scale)
K
Alveolar Septae (AS)
Blood vessel inner (intima)
Blood vessel outer (adventitia)
Airway (panCK-)
Airway (panCK+)
Infiltrate (CD45-)
Infiltrate (CD45+)
TNC+ core
Perifocal R1 (panCK+)
Perifocal R2 (panCK+
Perifocal R3 (panCK+)
Pericocal R1 (panCK-)
Perifocal R2 (panCK-)
Perifocal R3 (panCK-)
Control
IPF
tSNE1
tSNE2
Reference cell types
Activated Fibrotic FBs
Adventitial FBs
Alveolar FBs
Alveolar Macrophages
AT1
Arteriole
AT2
B-cells
Basal
Basophils
Capillary
CD4+Tcells
CD8+Tcells
Goblet
Interstitial Macrophages
KRT5-/KRT17+
Lymphatic
Mast
Monocytes MDMs
Multiciliated
Myofibroblasts
NK/NKT
pDCs
Plasma
Proliferating Airway
Proliferating Myeloid
RASC
SMCs Pericytes
SPP1+ Macrophages
Transitional AT2
Tregs
Venous
L
Z-score
*: p < 0.05
**: p < 0.01
***: p < 0.001
Airway epithelial cells
Airway goblet cells
Alveolar macrophages
Ciliated cells
Clara cells
Ionocytes
Pulmonary ATI cells
Pulmonary AT II cells

**Supplementary Fig. S2**: **Anatomical niche-driven ROI selection and validation**. (**A**) Bar plot of ROIs captured for RNA (blue, n=1,257) and protein (orange, n=1,362) spatial profiling, broken down by annotated microenvironment. Percentages represent the proportion of total ROIs per modality. (**B**) Immunofluorescence images of example regions of interest selected for GeoMx spatial profiling, showing DAPI (nuclei), PanCK (green), TNC (red) and CD45 (yellow) staining. Left-hand images show immunofluorescence; right-hand images show selected ROIs overlaid. From top: alveolar septae; blood vessel separated into inner (intimal) and outer (adventitial) compartments; airway ROIs segmented into epithelial ($PanCK^+$) and stromal ($PanCK^-$) compartments; immune infiltrate ROIs segmented into immune ($CD45^+$) and stromal ($CD45^-$) compartments; $TNC^+$ areas of active fibrogenesis surrounded by three concentric perifocal regions (R1-R3), each 90 μm in thickness, extending ~270 μm from the $TNC^+$ boundary. Perifocal regions are further segmented into epithelial ($PanCK^+$) and stromal ($PanCK^-$) compartments. (**C**) Laser-capture microdissection RNA-seq (n=10 individual control and IPF donors) comparing *TNC* and *COL1A2* expression between control alveolar septae, IPF alveolar septae, and fibroblastic foci. Relative expression levels are calculated as counts per million reads (CPM). Data from GSE169500.[9] (**D**) GeoMx digital spatial profiling expression of *TNC*, *COL1A1* and *ACTA2*, across 66 labelled anatomical regions. Data from Eyres et al.[4] (**E-H**) Immunofluorescence images of an IPF tissue section showing (**E**) nuclear staining (DAPI, blue), (**F**) TNC (red), (**G**) α-smooth muscle actin (α-SMA, yellow), and (**H**) merged channels. White-bordered insets show full-tissue overviews; main panels show zoomed regions. White arrows denote area of $TNC^+$ morphologically consistent with a fibroblastic focus. Scale bar (H): 100 μm. (**I**) Representative staining-based segmentation masks showing $TNC^+$ core and $PanCK^-$ perifocal regions R1-R3 (top row), and $TNC^+$ core and $PanCK^+$ perifocal regions R1-R3 (bottom row). (**J**) Log-scaled expression of genes (*AGER*, *ACTA2*, *ELN*, *TNC*, *KRT17*, *PTPRC*, *SFTPC*, *MMP7*, *COL1A1*, *COL1A2*, *MMP2*, *POSTN*) across all sampled subregions; jitter points represent individual ROIs. (**K**) t-SNE plots of segregated subregion ROIs across control and IPF ROIS, and principal cell type assigned to each region. (**L**) Cell-type enrichment across anatomical subregions inferred via decoupler (univariate linear model) using the PanglaoDB lung reference. Colour bar represents normalised enrichment across subregions (z-score); statistical significance shown by median adjusted p-values (*$p<0.05$, **$p<0.01$, ***$p<0.001$).

Figure S3

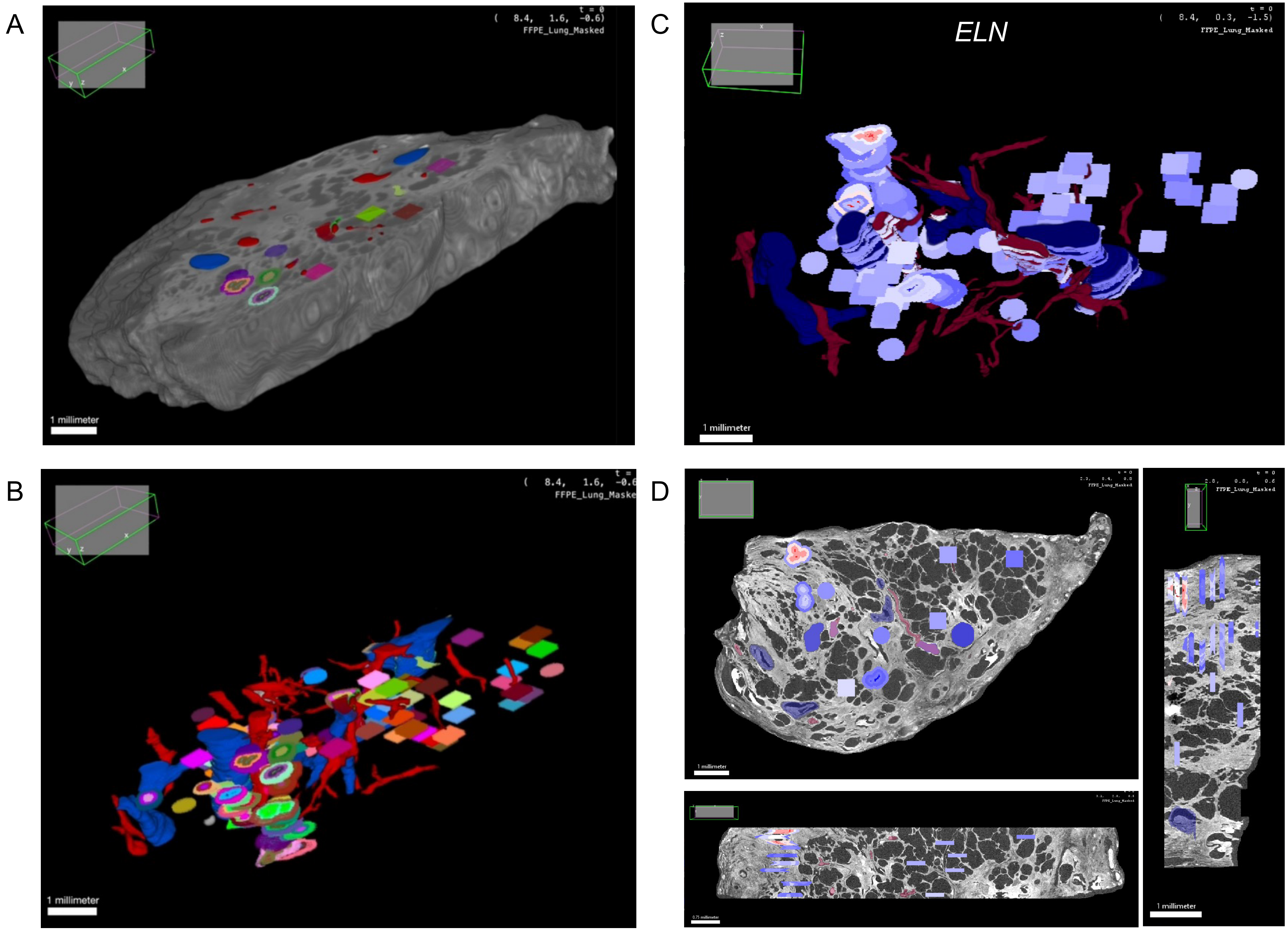
A
( 8.4, 1.6, -0.6)
FFPE_Lung_Masked
1 millimeter
B
1 millimeter
C
ELN
FFPE_Lung_Masked
1 millimeter
D
1 millimeter
1 millimeter

**Supplementary Fig. S3: Interactive 3D visualisation of spatially integrated molecular and structural data using the MoBIE plugin for Fiji. (A-D)** Screenshots from MoBIE, a Fiji plugin for sharing and exploring multimodal big-image data,[27] showing an IPF X-ray histology (XRH) volume with co-registered molecular and anatomical data.. **(A)** 3D volume rendering with surface-rendered volumes of interest (VOIs). **(B)** XRH data hidden, revealing 3D distributed VOIs and segmented vessels and airways. **(C)** 3D spatially resolved gene expression rendering: VOI-level expression of elastin (ELN), shown as an illustrative example, combined with segmented vessels and airways. Quantitative *ELN* expression across all sampled subregions is shown in the multi-gene expression overview in Fig. S2J. **(D)** Orthogonal cross-sections (XY, XZ, YZ planes) through the same XRH tissue volume, with VOIs coloured by ELN-expression overlaid overlaid. Insets in each panel indicate the current viewport orientation within the MoBIE interface.

### Figure S4

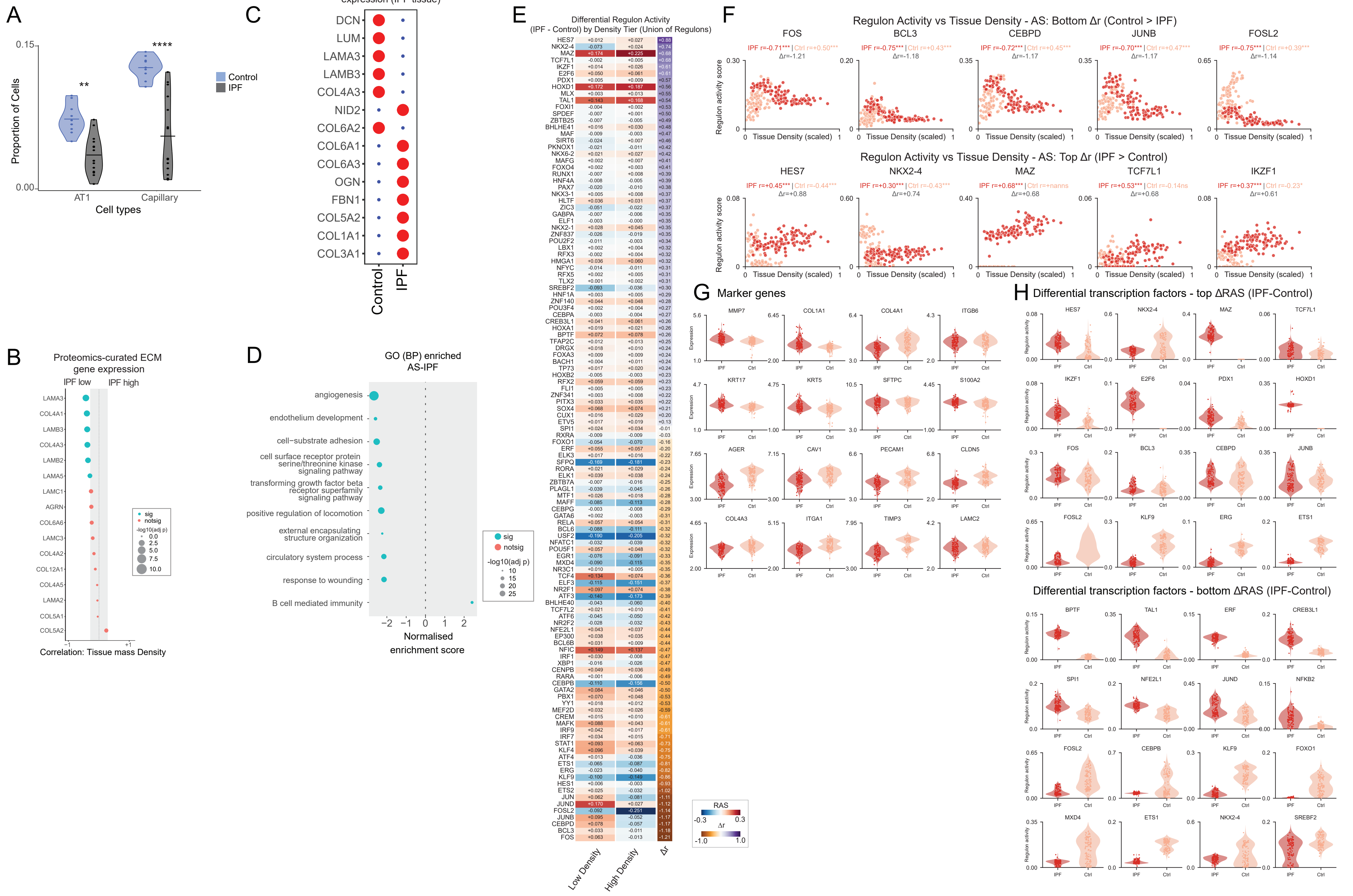

A
Proportion of Cells
Cell types
AT1
Capillary
Control
IPF
B
Proteomics-curated ECM gene expression
IPF low
IPF high
Correlation: Tissue mass Density
C
Proteomic-derived gene expression (IPF tissue)
DCN
LUM
LAMA3
LAMB3
COL4A3
NID2
COL6A2
COL6A1
COL6A3
OGN
FBN1
COL5A2
COL1A1
COL3A1
Control
IPF
D
GO (BP) enriched AS-IPF
angiogenesis
endothelium development
cell−substrate adhesion
cell surface receptor protein serine/threonine kinase signaling pathway
transforming growth factor beta receptor superfamily signaling pathway
positive regulation of locomotion
external encapsulating structure organization
circulatory system process
response to wounding
B cell mediated immunity
Normalised enrichment score
sig
notsig
-log10(adj p)
E
Differential Regulon Activity
(IPF - Control) by Density Tier (Union of Regulons)
Low Density
High Density
Δr
RAS
F
Regulon Activity vs Tissue Density - AS: Bottom Δr (Control > IPF)
FOS
BCL3
CEBPD
JUNB
FOSL2
Regulon Activity vs Tissue Density - AS: Top Δr (IPF > Control)
HES7
NKX2-4
MAZ
TCF7L1
IKZF1
Regulon activity score
Tissue Density (scaled)
G
Marker genes
Expression
H
Differential transcription factors - top ΔRAS (IPF-Control)
Differential transcription factors - bottom ΔRAS (IPF-Control)
Regulon activity

**Supplementary Fig. S4: Tissue mass density and disease-driven changes in matrix-associated septal gene modules**. (**A**) Cell-type proportion violins for AT1 and capillary endothelial cells, comparing control (or low-density) and IPF (or high-density) alveolar septae ROIs; $^{**}p < 0.01$, $^{****}p < 0.0001$ (t-test). (**B**) Correlation of gene expression with tissue mass density in IPF tissue, using a curated ECM gene list identified from an independent proteomic dataset of insoluble protein fractions identified as differentially abundant (IPF high or IPF low) in IPF compared to control tissue.[31] (**C**) Binary plot showing up- or down-regulation of genes derived from an independent quantitative human lung proteomic dataset of ECM proteins up or downregulated in IPF compared to control tissue [30]. (**D**) Top 10 significantly enriched Gene Ontology Biological Process terms from gene-set enrichment analysis comparing IPF vs control alveolar septae; bubble size represents -log10 adjusted p-value, colour indicates significance/direction. (**E**) Heatmap of regulon activity scores correlated with tissue density across IPF and control alveolar septae for all regulons; columns show mean differential regulon activity scores (ΔRAS, IPF-Control) and the rightmost column shows correlation difference with density ($\Delta r = r_{\_IPF} - r_{\_Control}$). (**F**) Scatter plots of regulon activity scores (y-axis) versus scaled tissue mass density (x-axis) for the top 10 differentially density-correlated regulons: top row shows the top 5 negative (Control>IPF); bottom row shows the top 5 positive (IPF>Control). Individual ROIs are shown as points coloured by group (IPF [colour], Control [colour]); annotations show group-specific Spearman correlations (r, Δr) and significance ($^{*}p < 0.05$, $^{**}p < 0.01$, $^{***}p < 0.001$). (**G**) Violin plots comparing log-scaled expression of region-specific marker genes between IPF and control alveolar septae ROIs. **(H)** Violin plots comparing mean regulon activity scores (RAS) for top- and bottom-ranked differentially active transcription factors between IPF and control alveolar septae ROIs.

### Figure S5

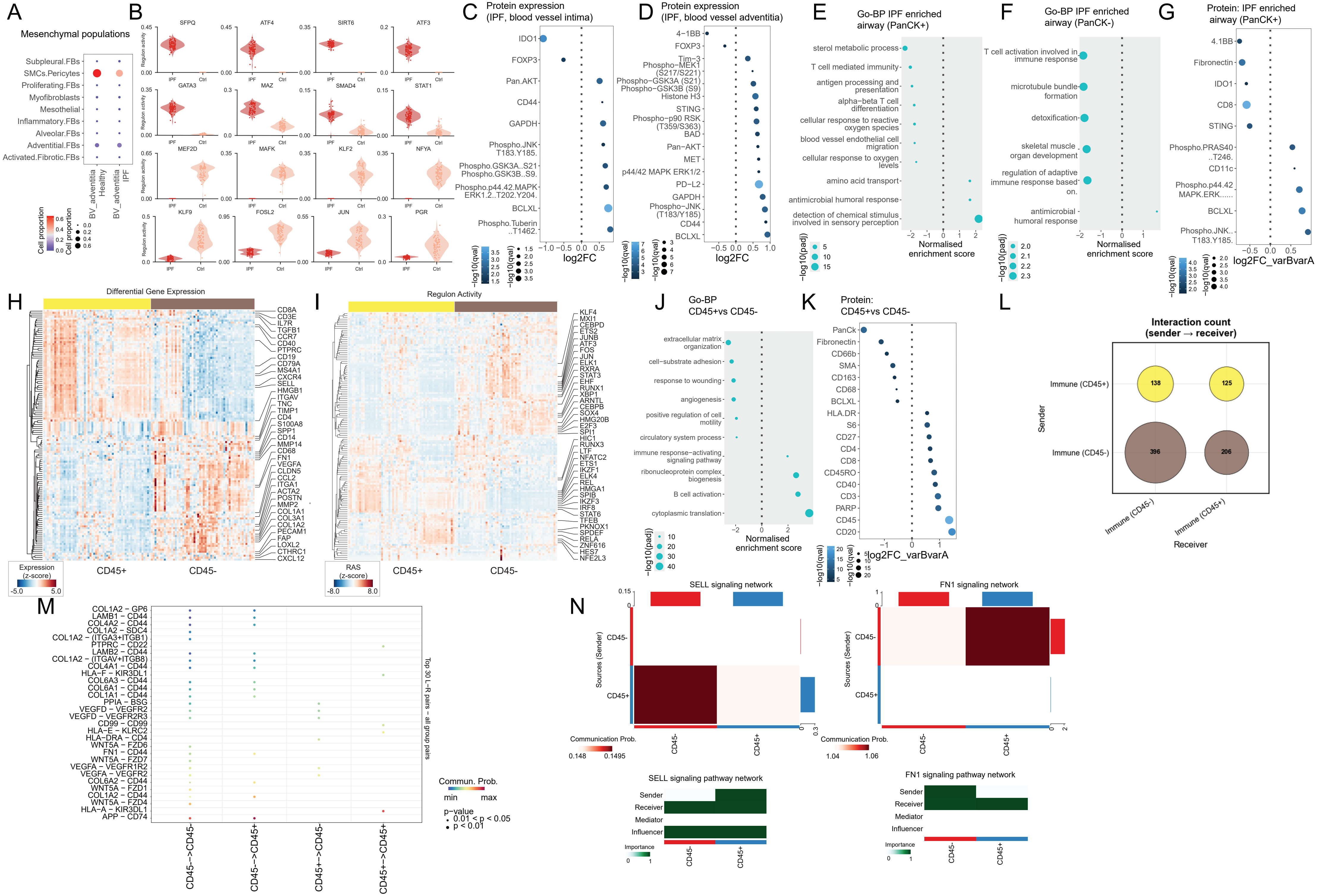

A
Mesenchymal populations
B
C
Protein expression (IPF, blood vessel intima)
D
Protein expression (IPF, blood vessel adventitia)
E
Go-BP IPF enriched airway (PanCK+)
F
Go-BP IPF enriched airway (PanCK-)
G
Protein: IPF enriched airway (PanCK+)
H
Differential Gene Expression
I
Regulon Activity
J
Go-BP CD45+vs CD45-
K
Protein: CD45+vs CD45-
L
Interaction count (sender → receiver)
M
N
SELL signaling network
FN1 signaling network
SELL signaling pathway network
FN1 signaling pathway network

**Supplementary Fig. S5: Altered molecular signatures in diverse pathogenic niches.** (**A**) Bubble plot showing mesenchymal cell-subtype proportions inferred by spatial deconvolution, comparing healthy and IPF blood-vessel adventitia compartments. (**B**) Violin plots comparing mean regulon activity scores (RAS) for top-ranked transcription factors between IPF and control blood-vessel ROIs. (**C**, **D**) Log-fold change/significance bubble plots showing differentially regulated proteins between IPF and control in inner (intimal, **C**) and outer (adventitial, **D**) blood-vessel compartments. (**E**, **F**) Normalised enrichment score/significance bubble plots showing top Gene Ontology Biological Process gene sets up- and down-regulated between IPF and control in PanCK$^{+}$ (**E**) and PanCK$^{-}$ (**F**) airway compartments. (**G**) Log-fold change/significance bubble plot showing differentially regulated proteins between IPF and control in PanCK$^{+}$ airway compartments. (**H**) Hierarchically clustered heatmap of the top differentially expressed RNA marker genes between CD45$^{+}$ and CD45$^{-}$ subregions within the immune-infiltrate compartment. (**I**) Heatmap of differential regulon activity (z-scored Regulon Activity Score, RAS) comparing CD45$^{+}$ and CD45$^{-}$ immune subregions. Colour bars in (**H**) and **(I)** indicate normalised expression and mean RAS (z-score). (**J**) Normalised enrichment score/significance bubble plot showing top Gene Ontology Biological Process gene sets up- and down-regulated between CD45$^{+}$ and CD45$^{-}$ subregions. (**K**) Log-fold change/significance bubble plot showing differentially regulated proteins between CD45$^{+}$ and CD45$^{-}$ subregions. (**L**) Bubble plot summarising the total number of significant ligand-receptor interactions between CD45$^{+}$ immune and CD45$^{-}$ parenchymal compartments, including both across-compartment and within-compartment (autocrine) signalling. Bubble area is proportional to number of significant interactions ($p < 0.05$). (**M**) Top 30 ligand-receptor pairs ranked by total communication probability. Each row highlights a ligand-receptor pair across four directed sender-to-receiver combinations. Bubble colour indicates communication probability; size reflects statistical significance ($p < 0.01$ or $0.01 \leq p < 0.05$). (**N**) Signalling role analysis for the SELL (left) and FN1 (right) pathways. Upper panels show pairwise communication probability between $CD45^{+}$ and $CD45^{-}$ compartments, with colour scaled to the observed probability range indicated in the colour bar; row bars (right) and column bars (top) indicate the summed outgoing and incoming communication strength per compartment respectively. Lower panels show network centrality scores quantifying the signalling role of each compartment as sender, receiver, mediator, and influencer, computed from the weighted directed communication network. Importance scores are normalised to a 0-1 scale (green colour bar) across $CD45^{-}$ and $CD45^{+}$ compartments.

### Figure S6

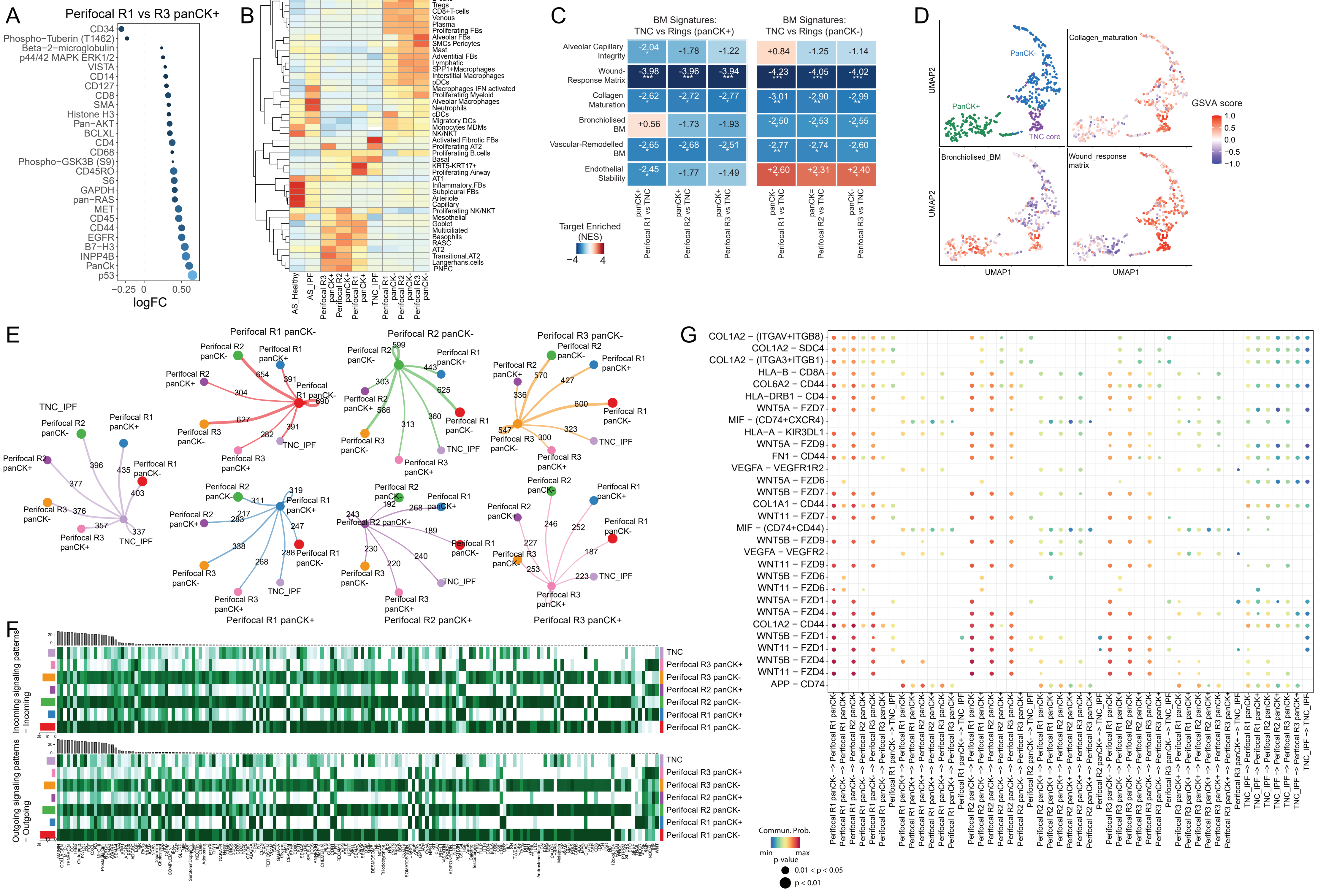

A
Perifocal R1 vs R3 panCK+
logFC
B
C
BM Signatures: TNC vs Rings (panCK+)
BM Signatures: TNC vs Rings (panCK-)
Alveolar Capillary Integrity
Wound-Response Matrix
Collagen Maturation
Bronchiolised BM
Vascular-Remodelled BM
Endothelial Stability
Target Enriched (NES)
D
PanCK-
PanCK+
TNC core
Collagen_maturation
Bronchiolised_BM
Wound_response matrix
GSVA score
UMAP1
UMAP2
E
Perifocal R1 panCK-
Perifocal R2 panCK-
Perifocal R3 panCK-
TNC_IPF
Perifocal R1 panCK+
Perifocal R2 panCK+
Perifocal R3 panCK+
F
Incoming signaling patterns – Incoming
Outgoing signaling patterns – Outgoing
G
Commun. Prob.
p-value
0.01 < p < 0.05
p < 0.01

**Supplementary Fig. S6: Perifocal cellular and molecular analyses.** (**A**) Log-fold change/significance bubble plot showing differentially abundant proteins between adjacent (Perifocal R1) and distal (Perifocal R3) PanCK$^+$ compartments. (**B**) Heatmap showing SpatialDecon-inferred cellular proportions in IPF and control alveolar septae, perifocal regions, and TNC$^+$ core compartments. (**C**) GSEA Normalised Enrichment Scores (NES) evaluating spatial shifts in basement-membrane gene signatures comparing the TNC$^+$ core to adjacent PanCK$^+$ (left heatmap) and PanCK$^-$ (right heatmap) perifocal regions. Colour gradient reflects normalised enrichment score; overlaid asterisks indicate significance (* FDR < 0.05, ** FDR < 0.01, *** FDR < 0.001). (**D**) UMAP projection of perifocal and TNC$^+$ core ROIs, shown as four panels coloured by anatomical class (PanCK$^+$ regions, PanCK$^-$ regions, TNC$^+$ core), collagen-maturation signature, bronchiolised basement-membrane signature, and wound-response matrix signature. (**E**) Compartment-centric circle plots showing outgoing ligand-receptor interactions originating from each perifocal compartment and the TNC$^+$ core; colour code matches Fig. 6**J**. Edge widths are proportional to the number of significant interactions, with numerical interaction counts indicated above edges. (**F**) Non-negative matrix factorisation (NMF) derived heatmaps of incoming (upper) and outgoing (lower) signalling patterns across all perifocal compartments and the TNC$^+$ core. Rows represent perifocal/core compartments, columns represent ligand-receptor signalling pathways, and colour intensity reflects NMF-derived contribution score (0 = white to maximum = dark green). (**G**) Top 30 ligand-receptor pairs ranked by total communication probability across perifocal regions and the TNC$^+$ core. Each row highlights a ligand-receptor pair across four directed sender-to-receiver combinations. Bubble colour indicates communication probability; size reflects statistical significance ($p < 0.01$ or $0.01 \leq p < 0.05$).

### Figure S7

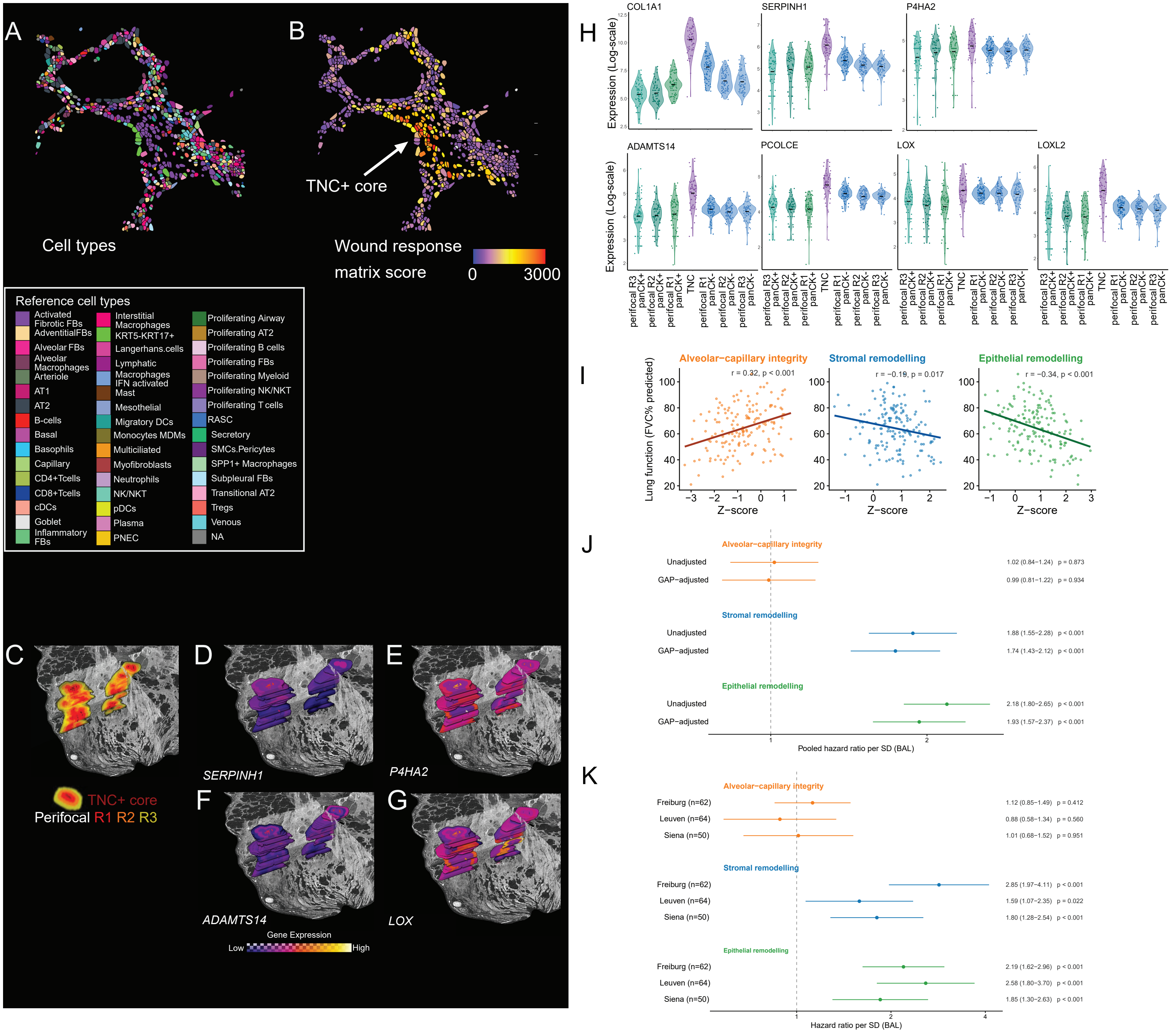

A
Cell types
B
TNC+ core
Wound response
matrix score
0
3000
Reference cell types
Activated Fibrotic FBs
AdventitialFBs
Alveolar FBs
Alveolar Macrophages
Arteriole
AT1
AT2
B-cells
Basal
Basophils
Capillary
CD4+Tcells
CD8+Tcells
cDCs
Goblet
Inflammatory FBs
Interstitial Macrophages
KRT5-KRT17+
Langerhans.cells
Lymphatic
Macrophages IFN activated
Mast
Mesothelial
Migratory DCs
Monocytes MDMs
Multiciliated
Myofibroblasts
Neutrophils
NK/NKT
pDCs
Plasma
PNEC
Proliferating Airway
Proliferating AT2
Proliferating B cells
Proliferating FBs
Proliferating Myeloid
Proliferating NK/NKT
Proliferating T cells
RASC
Secretory
SMCs.Pericytes
SPP1+ Macrophages
Subpleural FBs
Transitional AT2
Tregs
Venous
NA
C
D
E
TNC+ core
Perifocal R1 R2 R3
SERPINH1
P4HA2
F
G
ADAMTS14
LOX
Gene Expression
Low
High
H
COL1A1
SERPINH1
P4HA2
ADAMTS14
PCOLCE
LOX
LOXL2
Expression (Log-scale)
perifocal R3 panCK+
perifocal R2 panCK+
perifocal R1 panCK+
TNC
perifocal R1 panCK-
perifocal R2 panCK-
perifocal R3 panCK-
I
Alveolar−capillary integrity
Stromal remodelling
Epithelial remodelling
r = 0.32, p < 0.001
r = −0.19, p = 0.017
r = −0.34, p < 0.001
Lung function (FVC% predicted)
Z−score
J
Alveolar−capillary integrity
Unadjusted 1.02 (0.84−1.24) p = 0.873
GAP−adjusted 0.99 (0.81−1.22) p = 0.934
Stromal remodelling
Unadjusted 1.88 (1.55−2.28) p < 0.001
GAP−adjusted 1.74 (1.43−2.12) p < 0.001
Epithelial remodelling
Unadjusted 2.18 (1.80−2.65) p < 0.001
GAP−adjusted 1.93 (1.57−2.37) p < 0.001
Pooled hazard ratio per SD (BAL)
K
Alveolar−capillary integrity
Freiburg (n=62) 1.12 (0.85−1.49) p = 0.412
Leuven (n=64) 0.88 (0.58−1.34) p = 0.560
Siena (n=50) 1.01 (0.68−1.52) p = 0.951
Stromal remodelling
Freiburg (n=62) 2.85 (1.97−4.11) p < 0.001
Leuven (n=64) 1.59 (1.07−2.35) p = 0.022
Siena (n=50) 1.80 (1.28−2.54) p < 0.001
Epithelial remodelling
Freiburg (n=62) 2.19 (1.62−2.96) p < 0.001
Leuven (n=64) 2.58 (1.80−3.70) p < 0.001
Siena (n=50) 1.85 (1.30−2.63) p < 0.001
Hazard ratio per SD (BAL)

**Supplementary Fig. S7: Additional TNC⁺ core collagen maturation programme gene visualisation and association of atlas-derived signatures with clinical cohort outcomes.** (**A, B**) CosMx single-cell spatial maps showing annotated cell types (**A**) and wound-response matrix signature GSVA scores (**B**) within an IPF region containing a TNC+ core (white arrow). **(C-G)** Three-dimensional XRH rendering of $TNC^+$ cores and surrounding perifocal regions **(C)**, showing gene expression of *SERPINH1* (**D**), *P4HA2* (**E**), *ADAMSTS14* (**F**), and *LOX* **(G)** within $TNC^+$ cores and surrounding perifocal regions. **(H)** Log-scaled expression of genes (*COL1A1*, *SERPINH1*, *P4HA2*, *ADAMSTS14*, *PCOLCE*, *LOX*, *LOXL2*) across $TNC^+$ cores and perifocal subregions; jitter points represent individual ROIs. (**I-K**) Clinical correlation of atlas-derived spatial gene signatures . (**I**) Signature score (singScore, z) versus forced vital capacity (FVC, % predicted) in IPF lung tissue (LGRC/GSE47460, n = 157)[38]: alveolar–capillary integrity, perifocal stromal remodelling, and perifocal epithelial remodelling; Spearman r and p values inset. (**J**) Pooled hazard ratios for mortality per standard deviation of signature score in bronchoalveolar lavage (GSE70866, n = 176), shown unadjusted and adjusted for the GAP index. (**K**) Corresponding per-cohort hazard ratios (Freiburg, Leuven and Siena; per-cohort estimates are unadjusted). Points, hazard ratios (HR); bars, 95% confidence intervals; dashed line, HR = 1. Green, perifocal epithelial remodelling; blue, perifocal stromal remodelling; orange, alveolar-capillary integrity.

**Supplementary Table Legends**

**Table S1: GeoMx data:** Related to Fig 2 and S2. (**A**) Differentially expressed genes for each ROI type generated using EdgeR. Related to Fig. 2C.

**Table S2: Alveolar septae density-gene expression changes.** Related to Fig. 4 and S4. (**A**) Correlation of gene expression with tissue mass density. Related to Fig. 4D (**B**) Differential regulons across IPF and control septae, alongside their correlation to tissue density. Related to Fig. 4S. (**C**) All regulon activity scores correlated with tissue density. Related to Fig. S4E.

**Table S3: Vascular and immune changes in fibrosis.** Related to Fig. 5 and S5. (**A**) Gene set enrichment analysis results comparing Control vs IPF Inner (intimal) blood vessel regions. Related to Fig. 5D. (**B**) Gene set enrichment analysis results comparing Control vs IPF Outer (Adventitial) blood vessel regions. Related to Fig. 5E. **(C)** Correlation of PanCK$^+$ Perifocal Ring 3 region GSVA scores with geodesic distances from nearest blood vessel region. Related to Fig 5H. (**D**) Pathway directionality analysis quantifying the relative contribution of CD45$^+$ and CD45$^-$ compartments as signal senders across curated ligand-receptor pathways. Related to Fig. 5J. (**E, F**) Ligand receptor interactions between CD45$^+$ and CD45$^-$ subregions, with the sender population defined as CD45$^+$ (**E**) or CD45$^-$ (**F**). Related to Fig. 5K-L. (**G, H**) Gene set enrichment analysis results comparing Control vs IPF PanCK$^+$ (**G**) or PanCK$^-$ (**H**) Airway regions. Related to Fig S5E-F. (**I**) Total number of significant ligand-receptor interactions between CD45$^+$ immune and CD45$^-$ parenchymal compartments, including both across-compartment and within-compartment (autocrine) signalling. Related to Fig S5L. (**J**) Ligand receptor pairs and communication probability between CD45$^+$ immune and CD45$^-$

parenchymal compartments. Related to Fig S5M. (**K**) Ligand-receptor pairs ranked by total communication probability. Related to Fig S5M.

**Table S4: Perifocal R1-R3 compartment analyses.** Related to Fig. 6 and S6. (**A**) Differentially expressed genes between Perifocal Ring 1 and Ring 3 for both $PanCK^{+}$ and $PanCK^{-}$ compartments. Related to Fig. 6B. (**B**) Mean regulon activity scores for $TNC^{+}$ core and perifocal regions. Related to Fig 6C. (**C**) SpatialDecon calculated cell proportions for different ROI types. Related to Fig 6D,E,L. (**D**) Ligand-receptor interactions between perifocal compartments and $TNC^{+}$ core. Related to Fig 6J. (**E**) Pathway sender interactions across $panCK^{-}$ stromal, $panCK^{+}$ epithelial regions and $TNC^{+}$ core. Related to Fig. 6K. (**F**) Differentially abundant proteins between perifocal $panCK^{+}$ ring 1 and ring 3. (**G**) Protein Differential abundance analysis comparing $TNC^{+}$ core-adjacent to distal perifocal $panCK^{-}$ rings. Related to Fig. S6A. (**H**) Non-negative matrix factorisation (NMF) scores of incoming and outgoing signalling patterns across all perifocal compartments and the $TNC^{+}$ core. Related to Fig S6F. (**I**) Ligand-receptor pairs ranked by total communication probability across perifocal regions and the $TNC^{+}$ core. Related to Fig. S6G.

**Table S5: Clinical cohort analyses.** Related to Fig. 6 and S7 (**A**) Gene sets used to calculate singScore gene expression signatures. (**B**) Enrichment of spatial gene signatures among circulating IPF survival proteins (plasma proteome[40]). For each signature: number of genes, number measured on the Olink panel, overlap with combined-cohort validated prognostic proteins (and the risk-direction subset), and hypergeometric enrichment P value (overall survival-associated protein and adverse-outcome (risk) proteins only). Related to Fig 6T. (**C**) Signature genes that are present among circulating IPF survival proteins. Combined-cohort hazard ratios for the plasma proteins overlapping each signature; direction indicates association with increased (risk) or decreased (protective) risk of death/transplant. Related to Fig 6T.

**Table S6: Additional information.** (**A**) Demographic data of participants in this study (**B**) Summary of total and retained regulons per compartment after quality filtering. (**C**) Regulon Activity Scores (RAS) range per compartment. (**D**) Curated extracellular matrix gene sets. (**E**) Selected signalling pathways per spatial ROI subset for directional communication analysis.
